# Pheromone circuits and transcriptional cascades modulating transcriptional and chromatin states in the *Drosophila* central brain with social experience

**DOI:** 10.1101/2025.09.12.675922

**Authors:** Chengcheng Du, Jesús Emiliano Sotelo Fonseca, Lanling Jia, Shayna Scott, Sumie Okuwa, Sare Koruk, Corbin D. Jones, Pelin C. Volkan

**Affiliations:** Department of Biology, Duke University, Durham, NC 27708, USA; Department of Neurobiology, Duke University Medical Center, Durham, NC 27710, USA; Department of Biology, University of North Carolina at Chapel Hill, Chapel Hill, NC 27599, USA

## Abstract

Social experience significantly influences the behavioral and physiological responses of animals, including humans. In many animals, social isolation increases aggression, courtship, locomotion, and feeding while disrupting sleep. This occurs when peripheral neurons detect social signals, such as pheromones, which activate decision-making circuits in the brain. However, the molecular and circuit mechanisms of how chronic social isolation or enrichment alter gene expression and affect neuronal function and behavior remain unclear. In this study, we examined how transcription patterns and chromatin marks in male *Drosophila* brains change in response to social experience, and the effect of pheromone circuits and transcription factors involved in social circuit function. We focused on pheromone receptors Or47b and Or67d, as well as transcription factors Fru^M^ and Dsx^M^. Our findings suggest that social experience affects multiple genes in the central brain. Disrupting Or47b, Or67d, Fru^M^, and Dsx^M^ function moderated the transcriptional responses through antagonistic interactions. Specifically, Or47b circuits predominantly mediated transcriptional responses to social isolation through Dsx^M^ function, while Or67d and Fru^M^ regulated responses to group housing. Notably, mutants of *fru^M^* and *dsx^M^* exhibited more extensive transcriptional changes in the brain than Or mutants, especially for Fru^M^/Dsx^M^ target genes. While social experience did not lead to detectable alterations in the overall chromatin profile in the whole brain, mutants of the four genes resulted in significant changes in the enrichment of H3K4me3 and RNA polymerase II (RNAPolII) compared to wild type. Furthermore, mutants in *fru^M^* and *dsx^M^* generally eliminated social experience-dependent changes in sleep and locomotion behaviors, whereas Or mutants exhibited more modest disruptions. Overall, our results uncover the pheromone circuits and transcriptional cascades in regulating molecular and behavioral responses to social experience.

## Introduction

Across species, from invertebrates to mammals, social experience profoundly shapes behavior and physiology (Anders, 1978; Hoffman et al., 1975; Li et al., 2021; Luciano and Lore, 1975; Makinodan et al., 2012; Matsumoto et al., 2005; McGowan et al., 2009; Sánchez et al., 1998; Stagkourakis et al., 2020; Ueda and Kidokoro, 2002; Wang et al., 2022). Isolation disrupts cognitive processes such as sleep, memory, learning, and attention, while promoting maladaptive behaviors including aggression and excessive feeding (Brown et al., 2017; Cacioppo and Hawkley, 2009; Cacioppo et al., 2006; Cohen, 2004; Liu et al., 2025; Zelikowsky et al., 2018). In humans, these disruptions are associated with increased vulnerability to neuropsychiatric and neurodegenerative disorders (Cacioppo et al., 2015; Cacioppo et al., 2006; Cohen, 2004). Heightened awareness of these outcomes has led to the classification of social isolation and loneliness as a public health epidemic (General, 2023; Jeste et al., 2020; Lin, 2023). Yet, the molecular and circuit-level mechanisms mediating these effects remain largely unresolved.

Social experience has been shown to influence gene regulation in the nervous systems, from insects to mammals (Guerrero et al., 2020; Li et al., 2021; Robinson et al., 2008; Tung et al., 2012; Zhao et al., 2020). *Drosophila melanogaster*, as an excellent behavioral model, has been used to study many social behaviors, as well as behaviors influenced by social experience (Deanhardt et al., 2023; Ganguly-Fitzgerald et al., 2006; Goncharova et al., 2016; Li et al., 2021; Liu et al., 2011; Pan and Baker, 2014; Sethi et al., 2019; Ueda and Kidokoro, 2002; Ueda and Wu, 2009; Zhao et al., 2024; Zhao et al., 2020). *Drosophila* detects social cues by the peripheral neurons tuned to social cues like pheromones, which activate the pheromone and social circuits in the central brain, where information is integrated, and behavioral decisions are made (Fig. 1A). In the case of chronic social enrichment or isolation, the activity state of neuronal circuits activated by social experience, or lack thereof, likely alters gene expression at the transcription levels and chromatin patterns, ultimately adaptively modifying circuit function in the brain and behavioral outputs (Agrawal et al., 2019; Agrawal et al., 2020; Cushing and Kramer, 2005; Deanhardt et al., 2023; Flavell and Greenberg, 2008; Glastad et al., 2020; McGowan et al., 2009; Pan and Baker, 2014; Siuda et al., 2014; West and Greenberg, 2011; Wu et al., 2019; Zhao et al., 2020). Previous research on the *Drosophila* peripheral olfactory system demonstrated that social experience and pheromone receptor signaling modify chromatin states and alter the expression of numerous neuromodulatory genes, in addition to changing neuronal responses (Deanhardt et al., 2023; Sethi et al., 2019; Zhao et al., 2020). These findings suggest a strong link between social signals, transcriptional and chromatin modulation, and neuronal responses within the peripheral nervous system. Although there is clear evidence that social experience can alter gene regulation in the peripheral nervous system, the changes in gene expression and chromatin states in the central brain in different social contexts and the circuits that mediate these changes remain unclear (Fig. 1A).

**Figure 1.**
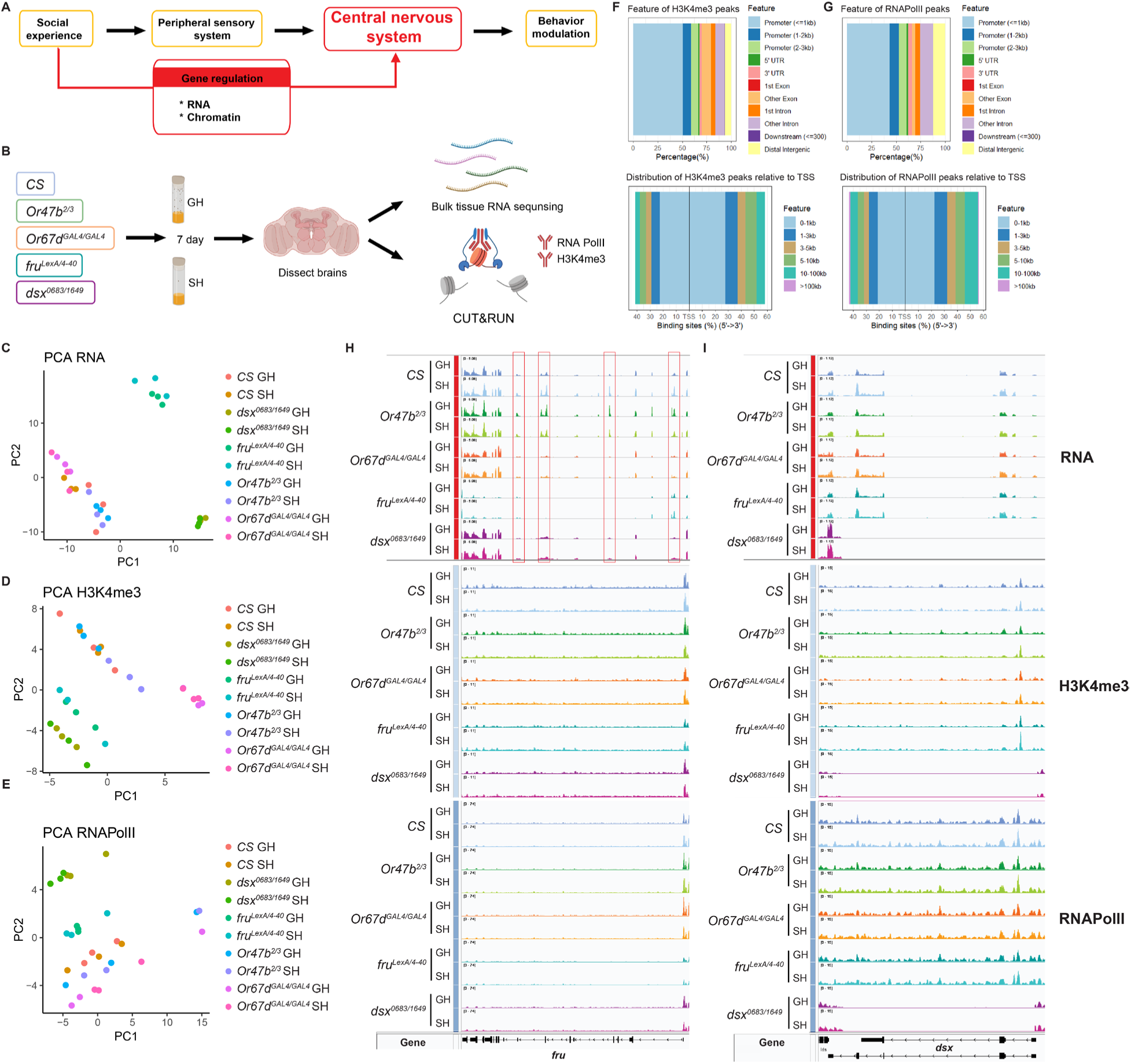
The effect of *Or47b, Or67d, fru^M^,* and *dsx^M^* mutants on transcriptional and chromatin states in male brains of GH and SH *Drosophila*. **(A)** Schematic for social signal transduction. **(B)** Brain RNA-seq and CUT&RUN workflow (Created with BioRender). **(C-E)** PCA analysis of transcriptional profiles **(C)**, and H3K4me3 **(D)** and RNAPolII enrichment chromatin profiles **(E)** from GH and SH wild-type *CS*, *Or47b^2/3^*, *Or67d^GAL4/GAL4^*, *fru^LexA/4–40^* and *dsx^0683/1649^* mutant male brains. **(F-G)** Feature distribution of H3K4me3 peaks **(F)** and RNAPolII peaks **(G)**. **(H-I)** IGV view of transcripts, H3K4me3 and RNAPolII distribution around *fru* (H) and *dsx* (I) locus among all genotypes (three biological replicates overlaid).

In *Drosophila*, two transcription factors (TFs), Fruitless (Fru) and Doublesex (Dsx), regulate the development and function of sex-specific social circuits and social behaviors (Anand et al., 2001; Burtis and Baker, 1989; Dauwalder, 2011; Demir and Dickson, 2005; Hildreth, 1965; Kimura et al., 2008; Nagoshi et al., 1988; Neville et al., 2014; Rideout et al., 2010; Ryner et al., 1996; Stockinger et al., 2005; Villella and Hall, 1996). The male isoform of Fru, Fru^M^, encodes a BTB zinc finger transcription factor involved in the sexual differentiation of neural circuits that control male behaviors (Ito et al., 1996; Ryner et al., 1996; Stockinger et al., 2005). In addition, *fru^M^* is necessary for mediating social experience-dependent changes in male competitive copulation behaviors (Sethi et al., 2019). We also recently reported that changes in *fru^M^* expression and function mediate social experience-dependent changes in courtship behaviors (Du et al., 2025). Fru was also shown to form a complex with the transcriptional cofactor Bonus (Bon), which recruits the chromatin regulators Histone deacetylase 1 (HDAC1) or Heterochromatin protein 1a (HP1a) to regulate gene expression underlying neuronal sexual differentiation (Ito et al., 2012). In contrast, Dsx belongs to the evolutionarily conserved DMRT (Doublesex and Mab-3 Related Transcription factor) gene family. Sex-specific isoforms of Dsx (Dsx^M^ and Dsx^F^) regulate sexual differentiation of both soma and the nervous system from invertebrates to vertebrates (Kopp, 2012; Murphy et al., 2007; Robinett et al., 2010). Perturbation of Dsx function changes courtship behavior in both males and females. Moreover, Dsx^M^ is necessary and sufficient for males to acquire experience-dependent courtship behavior in the absence of Fru^M^. Both Fru^M^ and Dsx^M^ bind DNA to regulate the expression of overlapping sets of genes, although some genes are specifically regulated by either Fru^M^ or Dsx (Clough et al., 2014; Neville et al., 2014).

Fru^M^ is expressed in interconnected neurons that label circuits for mediating social behaviors such as courtship and aggression (Cachero et al., 2010; Jai et al., 2010; Vrontou et al., 2006). Fru^M^ positive neural circuits include peripheral neurons that detect social cues to neural centers for sensory processing, integration, and decision-making, as well as the descending motor pathways responsible for executing the behaviors (Dauwalder, 2011; Jai et al., 2010; Kimura et al., 2008; Lee et al., 2000; Neville et al., 2014; Stockinger et al., 2005; von Philipsborn et al., 2014). In the olfactory system, two classes of pheromone-sensing olfactory receptor neurons (ORNs) that express pheromone receptors Or47b and Or67d are *fru^M^* positive (Jai et al., 2010; Stockinger et al., 2005). Or47b ORNs detect male and female pheromones and drive age and social experience-dependent male copulation advantage in a *fru^M^*-dependent manner (Dweck et al., 2015; Lin et al., 2016; Sethi et al., 2019). Or67d ORNs detect male-specific pheromone 11-*cis*-vaccenyl acetate (*cis*-VA) and function to inhibit courtship behaviors in males (Ha and Smith, 2006; Kurtovic et al., 2007).

Previous studies demonstrated that social isolation alters many behavioral and physiological responses of *Drosophila,* increasing aggression, courtship, locomotion, and feeding, and disrupting sleep (Deanhardt et al., 2023; Ganguly-Fitzgerald et al., 2006; Goncharova et al., 2016; Li et al., 2021; Liu et al., 2011; Pan and Baker, 2014; Sethi et al., 2019; Ueda and Kidokoro, 2002; Ueda and Wu, 2009; Zhao et al., 2024; Zhao et al., 2020). In a related study, we demonstrated that changes in courtship behaviors, resulting from social experiences, are associated with alterations in the expression and function of *fru^M^*, as well as specific circadian genes that are potential targets of Fru^M^ and/or Dsx^M^ (Du et al., 2025). Activating pheromone-sensing neurons through chronic exposure to social cues stimulates social circuits and engages the transcriptional functions of Fru^M^ and Dsx^M^ within these neurons.

Behavioral modulation in response to social experiences likely depends on detectors of social signals, such as pheromone receptors (like Or47b and Or67d), and regulators of social circuits like Fru^M^ and Dsx^M^. To test this hypothesis, we conducted a study exploring how transcriptional and chromatin patterns in the brain change when we functionally disrupt these four pathways using male mutants in *Or47b*, *Or67d*, *fru^M^*, and *dsx^M^* raised in monosexual groups or in isolation (Fig. 1B). We additionally examined the impact of these mutants on social modulation of behaviors. Our results suggest that: 1) Social experience leads to changes in numerous genes involved in various processes, from metabolism to behavioral regulation, neural development, and immunity. Loss of *Or47b*, *Or67d*, *fru^M^*, or *dsx^M^* normalized the transcriptional responses to social experience, with general antagonistic effects between Or47b and Or67d pheromone circuits, and between Fru^M^ and Dsx^M^ circuits. We found trends where Or47b pheromone circuits and Dsx^M^ circuits function together in mediating transcriptional responses to social isolation, while Or67d neuronal circuits and Fru^M^ circuits function together to drive transcriptional responses to group housing in the brain. 2) Under the same housing conditions, compared to Canton-S wild type (*CS*), *fru^M^* and *dsx^M^*mutant brains showed more extensive changes at the transcriptional level than those observed in *Or47b* or *Or67d* mutants. Fru^M^ and Dsx^M^ function as either transcriptional repressors or activators, influencing downstream target genes. 3) Despite transcriptional changes, at the chromatin level, social experience does not significantly alter the overall distribution of H3K4me3 or RNA polymerase II (RNAPolII) in the whole brain. However, compared to *CS*, the four mutants cause numerous changes to the enrichment of H3K4me3 or RNAPolII under the same social conditions. The transcriptional changes identified in different mutants are accompanied by modest concordant changes in H3K4me3 or RNAPolII enrichment. 4) Mutants in *fru^M^* and *dsx^M^*generally eliminate social experience-dependent changes in sleep and locomotion behaviors. In contrast, Or mutants exhibited more modest disruptions in social modulation of locomotion and courtship, with housing-specific defects observed in *Or47b* mutants. Our results identified pheromone circuits and transcriptional cascades that work with them in the modulation of molecular responses to social experience, which reprogram brain function and behaviors. These findings provide new insights into how social experiences affect molecular plasticity, laying the groundwork for future research on the molecular mechanisms linking social interactions to behavior and physiology.

## Results

### The contribution of pheromone receptors and social circuit regulators on transcriptional and chromatin responses to social experience in the brain

We recently reported differentially expressed genes (DEGs) between grouped and isolated *CS* male brains (Du et al., 2025). To explore the contribution of pheromone circuits and social circuit regulators in social modulation of gene expression, we analyzed transcriptional and chromatin profiles in *Or47b, Or67d, fru^M^*, and *dsx^M^* mutant brains from grouped or isolated males (Fig. 1A-B). We performed bulk RNA sequencing of brain tissues from group-housed (GH) and single-housed (SH) male *CS* wild type, and *Or47b^2/3^*, *Or67d^GAL4/GAL4^*, *fru^LexA/4-40^*, and *dsx^0683/1649^* mutants (Fig. 1B). To explore potential chromatin-based mechanisms for social experience-dependent gene regulation, we additionally conducted CUT&RUN (Cleavage Under Targets and Release Using Nuclease) and sequencing in male brains using the same genotypes and social conditions to assess the genome-wide enrichment of two active chromatin marks: H3K4me3 and RNAPolII (Fig. 1B).

We first examined the quality of the sequencing results. From the PCA results of both RNA seq and CUT&RUN, samples with the same genotypes are clustered together (Fig. 1C, D, and E). Both the transcription profiling and chromatin landscape of *fru^LexA/4-40^* and *dsx^0683/1649^* mutants appeared more distinct from the wild-type *CS*, while *Or47b^2/3^* and *Or67d^GAL4/GAL4^* mutants showed more modest changes compared to the wild type (Fig. 1C, D, and E). Regarding the distribution of H3K4me3 and RNAPolII peaks, over 50% of the peaks for both marks were found around the promoter regions throughout the genome (Fig. 1F and G).

Normalized counts of RNA transcripts of several housekeeping genes or control genes (*RpS29*, *Hmt4-20*, and *TSG101*) appeared consistent across all genotypes (Supplementary Fig. 1A-C). *fru^LexA/4-40^* mutants significantly decreased the expression of *fru* (Supplementary Fig. 1E). Within the overall brain, *fru* expression showed very little to no change with social experience in *CS* or mutants. Yet, similar to results from the peripheral olfactory system (Deanhardt et al., 2023), we found some *fru* exons to be differentially regulated in *CS* with social experience (Fig. 1H frames). The same exons were also decreased in *Or67d^GAL4/GAL4^* mutants and *dsx^0683/1649^* mutants (Fig. 1H frames). *fru^LexA/4-40^* males exhibited a reduction in H3K4me3 and RNAPolII enrichment at the *fru* locus (Fig. 1H). *dsx^0683/1649^* mutants also exhibited decreased *dsx* expression, as well as H3K4me3 and RNAPolII enrichment at the *dsx* locus (Fig. 1I; Supplementary Fig. 1F).

These results show that Fru^M^ and Dsx^M^ have a more dramatic impact on the transcriptional and chromatin responses to social experience, compared to the effect of Or47b and Or67d pheromone circuits.

### Or47b and Or67d pheromone circuits function antagonistically to influence transcriptional responses to social experience in the central brain

Compared to the SH *CS*, we found approximately 130 genes that are transcriptionally up- or downregulated in GH *CS* (Fig. 2A; Fig. 4A, B). These DEGs are involved in many biological processes, especially immunity, metabolism, receptor signaling, and behaviors (Fig. 2B; Fig. 3B; Supplementary Fig. 4A; Supplementary Table 1). We first examined how pheromone circuits drive transcription responses to social experience. Both *Or47b^2/3^* and *Or67d^GAL4/GAL4^* mutants showed a significant reduction in the number of DEGs between GH and SH (Fig. 4A-C). *Or47b^2/3^* mutants predominantly disrupted DEG patterns in response to social isolation (Fig. 2A). In contrast, in *Or67d^GAL4/GAL4^* mutants, DEG patterns in response to group housing were disrupted (Fig. 2A). We also compared the Gene Ontology (GO) terms of DEGs in GH/SH comparisons of *CS*, *Or47b^2/3^* and *Or67d^GAL4/GAL4^*mutants to determine if the two pheromone circuits modulate different biological processes and molecular functions in response to social experience (Fig. 2B). For example, DEGs related to immune functions in *CS* male brains were no longer DEGs in *Or67d^GAL4/GAL4^* mutant brains (Fig. 2A, B). This suggests that Or67d circuits may contribute to the transcriptional differences in immune genes between grouped and isolated wild-type male brains. In *Or47b^2/3^*mutants, on the other hand, most specific changes are observed in GO terms for locomotion and hormone activity (Fig. 2A, B).

**Figure 2.**
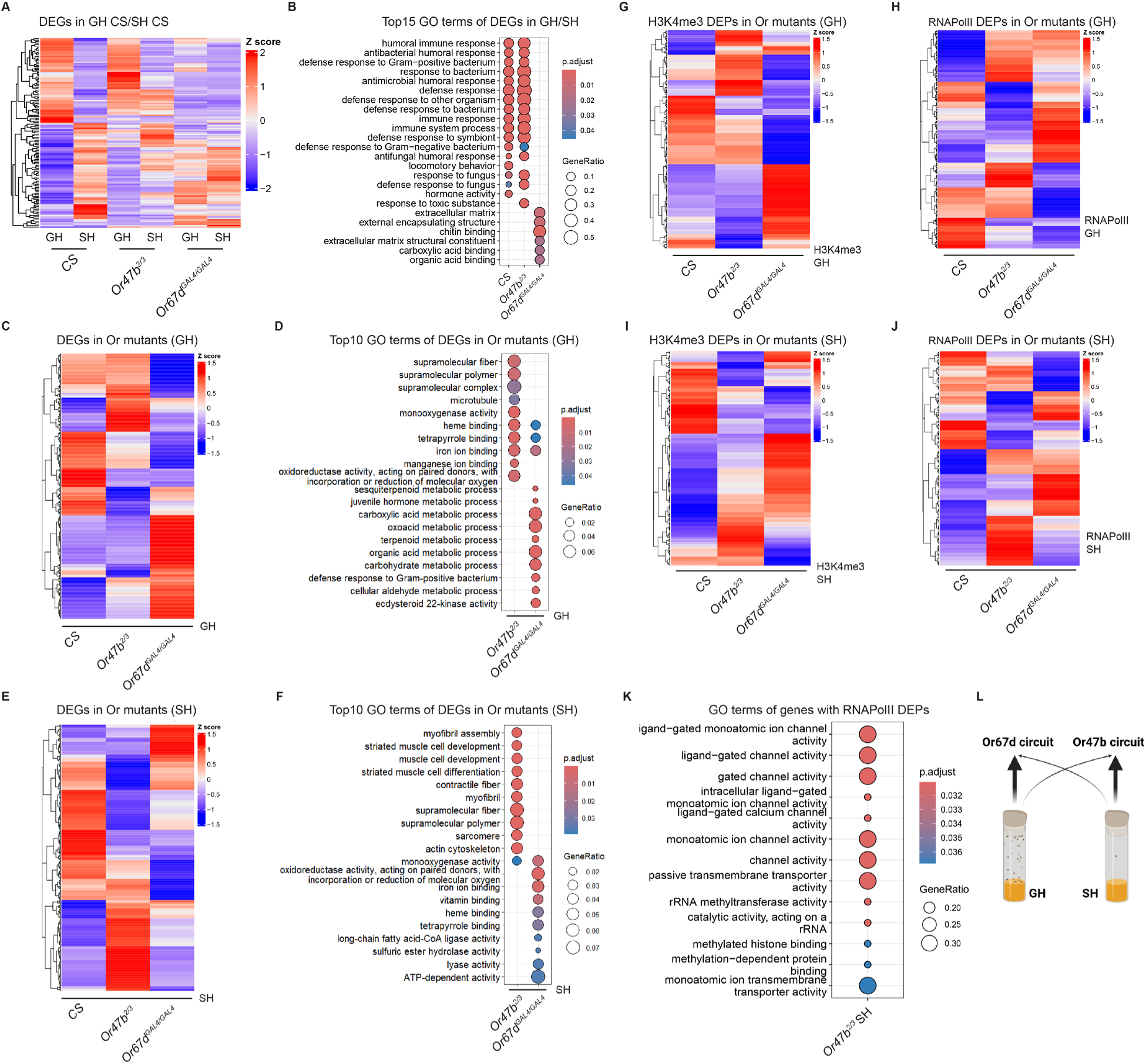
The effect of Or47b and Or67d pheromone circuits on transcriptional and chromatin responses to social experience in the central brain. **(A)** Heatmaps displaying Z-scores of the median of normalized transcript counts for DEGs identified in the male brains from the GH *CS*/SH *CS* comparison across *CS* and two Or mutants, *Or47b^2/3^* and *Or67d^GAL4/GAL4^*(adjusted p-value < 0.05, |log₂FC| > 0.1). **(B)** Top 15 most significantly enriched GO terms for DEGs of GH/SH comparisons of *CS* and two Or mutants, respectively (q-value < 0.05). **(C, E)** Heatmaps displaying Z-scores of the median of normalized transcript counts for DEGs identified from the Or mutants/*CS* comparison in GH **(C)** and SH **(E)** (adjusted P-value < 0.05, |log₂FC| > 0.1). **(D, F)** Top 10 most significantly enriched GO terms for DEGs of Or mutants/*CS* comparison in GH **(D)** and SH **(F)** (q-value < 0.05). **(G-H)** Heatmaps displaying Z-scores of the median of normalized enrichment levels of H3K4me3 **(G)** or RNAPolII **(H)** DEPs identified from Or mutants/*CS* comparison in GH brains (adjusted P-value < 0.01). **(I-J)** Heatmaps displaying Z-scores of the median of normalized enrichment levels of H3K4me3 **(I)** or RNAPolII **(J)** DEPs identified from Or mutants/*CS* comparison in SH brains (adjusted P-value < 0.01). **(K)** GO terms for genes with RNAPolII DEPs of *Or47b^2/3^* mutants/*CS* comparison in SH brains (q-value < 0.05). **(L)** A working model depicting the roles of Or47b and Or67d pheromone circuits in mediating transcriptional response to social experience in the central brain.

**Figure 3.**
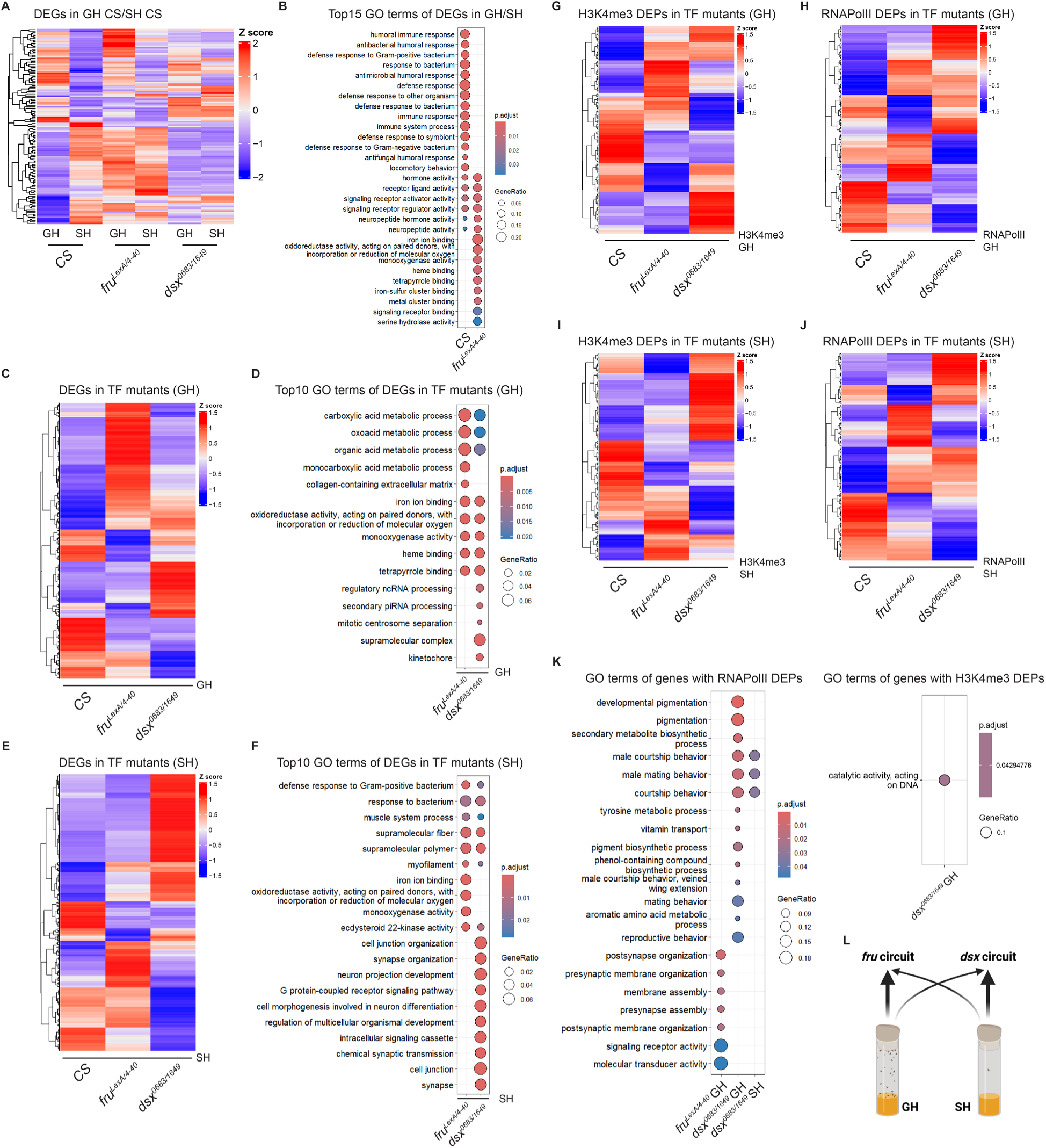
The effect of transcription factors Fru^M^ and Dsx^M^ on transcriptional and chromatin responses to social experience in the central brain. **(A)** Heatmaps displaying Z-scores of the median of normalized transcript counts for DEGs identified in the male brains from the GH *CS*/SH *CS* comparison across *CS,* and mutants in two transcription factors (TFs), *fru^LexA/4-40^*, and *dsx^0683/1649^* (adjusted p-value < 0.05, |log₂FC| > 0.1). **(B)** Top 15 most significantly enriched GO terms for DEGs of GH/SH comparisons of *CS* and *fru^LexA/4-40^*mutant, respectively (q-value < 0.05). **(C, E)** Heatmaps displaying Z-scores of the median of normalized transcript counts for DEGs identified from TF mutants/*CS* comparison in GH **(C)** and SH **(E)** (adjusted P-value < 0.05, |log₂FC| > 0.1). **(D, F)** Top 10 most significantly enriched GO terms for DEGs of TF mutants/*CS* comparison in GH **(D)** and SH **(F)** (q-value < 0.05). **(G-H)** Heatmaps displaying Z-scores of the median of normalized enrichment levels of H3K4me3 **(G)** or RNAPolII **(H)** DEPs identified from TF mutants/*CS* comparison in GH brains (adjusted P-value < 0.01). **(I-J)** Heatmaps displaying Z-scores of the median of normalized enrichment levels of H3K4me3 **(I)** or RNAPolII **(J)** DEPs identified from TF mutants/*CS* comparison in SH brains (adjusted P-value < 0.01). **(K)** GO terms for genes with H3K4me3 or RNAPolII DEPs of TF mutants/*CS* comparison in either GH or SH condition (q-value < 0.05). **(L)** A working model displaying the roles of Fru^M^ and Dsx^M^ circuits in mediating transcriptional response to social experience in the central brain.

**Figure 4:**
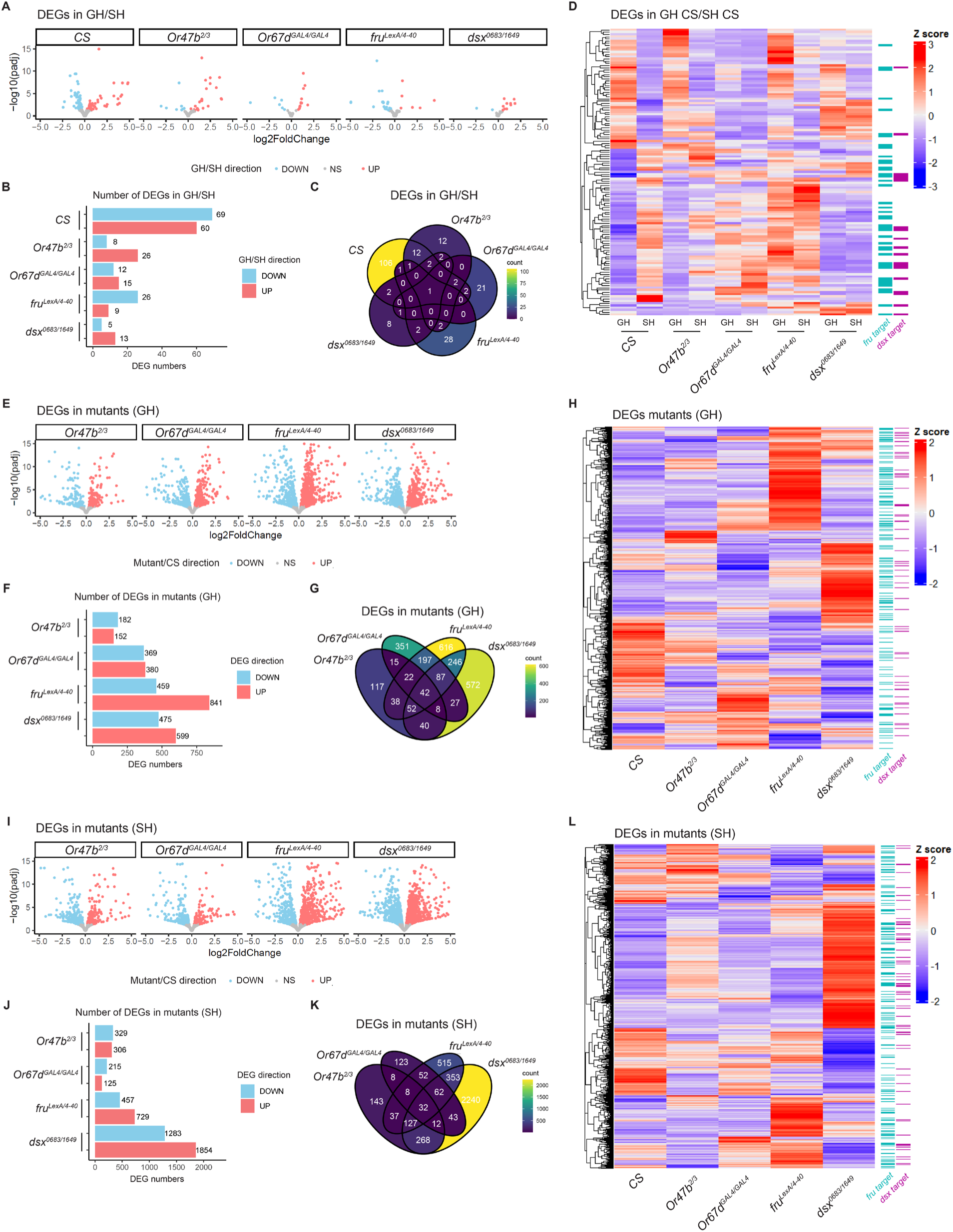
Comparisons of the transcriptional effects of Or and TF mutants in male fly brains. **(A)** Volcano plots displaying log_2_FoldChange (GH/SH) and -log10(p-adj) of genes in each genotype. DEGs were labeled in red (higher in GH) or blue (lower in GH). **(B)** Numbers of DEGs in GH/SH comparisons for each genotype (adjusted p-value < 0.05, |log₂FC| > 0.1). **(C)** Venn diagram comparing DEGs from GH/SH comparisons for each genotype. **(D)** Heatmaps displaying Z-scores of the median of normalized transcript counts for DEGs from the GH *CS*/SH *CS* comparison across *CS*, Or mutants, and TF mutants (adjusted p-value < 0.05, |log₂FC| > 0.1). Putative Fru^M^ or Dsx^M^ target genes are labeled on the right. **(E, I)** Volcano plots displaying log_2_FoldChange (mutant/*CS*) and -log10(p-adj) of genes in each genotype under GH **(E)** or SH **(I)** condition. DEGs up-and down-regulated in mutants are labeled in red and blue, respectively. **(F, J)** Numbers of DEGs in mutant/*CS* comparisons for each genotype under GH **(F)** or SH **(J)** condition (adjusted p-value < 0.05, |log₂FC| > 0.1). **(G, K)** Venn diagram comparing DEGs from mutant/*CS* comparisons for each genotype under GH **(G)** or SH **(K)** conditions. **(H, L)** Heatmaps displaying Z-scores of the median of normalized transcript counts for all DEGs from any of the mutant/*CS* comparison under either GH **(H)** or SH **(L)** condition (adjusted p-value < 0.05, |log₂FC| > 0.1). Putative Fru^M^ or Dsx^M^ targets are labeled on the right.

We next investigated how chromatin states changed in response to different social experiences. The overall distribution of either H3K4me3 or RNAPolII did not show any significant change between the GH/SH housing comparison in any genetic background (Fig. 6D), including *CS*, which had the most DEGs. Interestingly, multiple papers over the past two decades indicate that this is a relatively common observation (Agrawal et al., 2019; Brovkina et al., 2021; Kiani et al., 2022). This discrepancy, where DEGs are present without accompanying changes in chromatin marks, may be due to technical limitations in detecting chromatin level changes that occur only in specific cell populations within the brain. These localized changes might be masked when analyzing chromatin at the global level across the entire brain. Furthermore, the hundreds to thousands of RNA transcripts produced from a single gene in the genome— typically with only two copies in most cells—could also explain the absence of a significant change in chromatin enrichment for H3K4me3 or RNAPolII around DEGs under varying social conditions.

The results indicate that Or67d circuits primarily influence transcriptional changes in response to group housing. In contrast, Or47b circuits mainly drive transcriptional responses to social isolation, although their impact on the social modulation of transcriptional profiles in the brain is relatively weaker (Fig. 2L). These data highlight the differences and opposing functions of the two pheromone circuits in implementing the effects of social experience on gene expression throughout the brain.

### The effect of Or47b and Or67d pheromone circuits on gene expression and chromatin state in the brain

In addition to investigating the contribution of pheromone circuits in mediating social modulation of transcription, we next examined how pheromone circuits affected overall transcription under the same social condition. When compared to *CS* under both GH and SH conditions, the *Or47b^2/3^*mutant changed the expression of many supramolecular and cytoskeletal-related genes (Fig. 2C-F; Supplementary Fig. 5). *Or67d^GAL4/GAL4^*mutant, on the other hand, altered the expression of various metabolic processes and defense responses under GH conditions (Fig. 2C-D; Supplementary Fig. 5). Under SH conditions, the *Or67d^GAL4/GAL4^* mutant mainly altered the expression of genes in molecular binding and metabolic processes (Fig. 2E-F; Supplementary Fig. 5). Overall, the effect of disrupting Or67d and Or47b function on transcriptional profiles in both GH and SH brains was specific to each pheromone circuit, with little overlap between the mutants (Fig. 2C-F, Fig. 4G and K). These results agree with specific and overlapping roles for each pheromone circuit in mediating distinct cellular and transcriptional responses in the central brain.

We next investigated how chromatin states change when different pheromone circuits are disrupted. We did not find significant differentially enriched peaks (DEPs) for H3K4me3 or RNAPolII in male brains when comparing GH/SH housing conditions in any of the genotypes (Fig. 6D). However, loss of *Or47b* or *Or67d* exhibited DEPs for H3K4me3 and RNAPolII when compared to *CS* under either GH or SH conditions (Fig. 2G-K). The effect of Or47b or Or67d circuits on H3K4me3 and RNAPolII enrichment compared to wild type in both GH and SH brains was generally specific (Fig. 2G-J). A small portion of DEPs were altered in the same direction in both mutants, yet to different extents (Fig. 2G-J). To determine the genes with DEPs, we linked the DEPs to the nearest genes (Fig. 2G-K). Next, we performed GO term analysis on the genes with DEPs in *Or47b^2/3^* and *Or67d^GAL4/GAL4^* mutants compared to wild type in either social condition (Fig. 2K). We found that *Or47b^2/3^*mutants altered the RNAPolII distribution around genes involved in channel activity-related processes under the SH condition but not in the GH condition (Fig. 2K). Genes with RNAPolII or H3K4me3 DEPs in *Or67d^GAL4/GAL4^* mutants were not enriched for any GO terms.

We next extended the analysis to determine if the changes in DEPs were concordant with the changes in DEGs. We compared DEGs to genes with DEPs that changed in the same direction in Or mutants (Supplementary Fig. 2). The number of genes with DEPs was much less than that of DEGs. We identified a few genes exhibiting concordant changes in transcription and enrichment patterns for either active mark. The genes with concordant changes in *Or47b^2/3^* mutants are involved in various metabolic, catabolic, and intracellular transport processes, and while in *Or67d^GAL4/GAL4^* mutants, they are involved in enzyme activity, DNA, and chromosome regulation (Supplementary Fig. 2E-F). These results suggest that Or47b and Or67d ORN circuits reprogram chromatin states and transcription around unique sets of genes in the brain, highlighting potential differences in cellular responses driven by different pheromone circuits.

### Fru^M^ and Dsx^M^ function antagonistically to influence transcriptional responses to social experience in the central brain

The effect of Fru^M^ or Dsx^M^ on gene regulation in the nervous system, or their modes of transcriptional regulation, remains unclear. We performed a similar analysis on *fru^LexA/4-40^* and *dsx^0683/1649^* mutant brains. First, we investigated whether the function of Fru^M^ or Dsx^M^ is required to drive transcriptional changes to social experience observed in the wild type using *fru^LexA/4-40^* and *dsx^0683/1649^*mutant males that are either grouped or isolated. The expression pattern of DEGs in GH/SH comparisons in *CS* brains showed that Fru^M^ and Dsx^M^ have opposing roles in mediating gene expression with social experience. In *fru^LexA/4-40^*or *dsx^0683/1649^* mutants, the transcriptional differences between GH and SH are largely eliminated, albeit in different ways (Fig.3A; Fig.4A-D). Specifically, the transcriptional profile in both GH and SH *fru^LexA/4-40^* mutant brains appeared similar to that of SH *CS* brains (Fig.3A), suggesting that Fru^M^ contributes to mediating the transcriptional responses to GH. This pattern was opposite in *dsx^0683/1649^*mutant brains, where, for many genes, transcriptional responses to isolation appeared more significantly disrupted compared to GH (Fig.3A). This result indicates that, in contrast to Fru^M^, Dsx^M^ plays a significant role in mediating the transcriptional responses to SH. Fru^M^ or Dsx^M^ were previously shown to bind to the regulatory elements of many of the DEGs (Fig. 4D) (Clough et al., 2014; Neville et al., 2014), suggesting that the transcriptional defects observed are likely a direct downstream effect of the loss of Fru^M^ and Dsx^M^ function. The fact that some of these DEGs were upregulated and others were downregulated in *fru^LexA/4-40^* and *dsx^0683/1649^*mutants also reveals that both transcription factors (TFs) can function as transcriptional repressors or activators in a gene-specific manner (Fig. 4D, H, L; Supplementary Fig. 6). Alternatively, Fru and Dsx proteins can potentially form different homomeric or heteromeric complexes, which determine repressor or activator functions on target genes as well as themselves, and mutations in either *fru* or *dsx* can alter the stoichiometry of these complexes, determining their effects on transcription.

The total number of DEGs between GH and SH male brains is significantly lower in *fru^LexA/4-40^* or *dsx^0683/1649^*mutants compared to that of wild-type male brains (Fig. 4A-C). A number of GO terms related to metabolism and behavior from DEGs responding to social experience in *CS* were absent in *fru^LexA/4-40^*mutants (Fig.3A-B; Supplementary Table 1). Yet, the DEGs identified between GH and SH in *fru^LexA/4-40^* mutants are enriched in GO terms such as various receptor activities and binding processes (Fig. 3B; Supplementary Fig. 4). The DEGs between grouped and isolated *dsx^0683/1649^*mutants were not enriched in any GO terms but rather in defense response and cell communication Reactome or KEGG terms (Fig.3B; Supplementary Fig. 4). In addition to the normalization of transcription in response to social experience in *fru^LexA/4-40^* or *dsx^0683/1649^* mutants, the distribution of two active chromatin marks, H3K4me3 and RNAPolII, did not show any significant changes between housing conditions in either mutant (Fig. 6D). Overall, these data reveal opposing modes of regulation for Fru^M^ and Dsx^M^ in mediating transcriptional responses to grouping and isolation in the brain, respectively (Fig. 3L).

### The effect of Fru^M^ and Dsx^M^ function on gene expression and chromatin state in the brain

Next, we analyzed changes in transcription in *fru^LexA/4-40^*and *dsx^0683/1649^* mutant male brains compared to the wild type under the same housing conditions (Fig. 3C-F), which can include genes directly regulated by Fru^M^ and Dsx^M^. DEGs in GH conditions from *fru^LexA/4-40^* mutant/*CS* or *dsx^0683/1649^* mutant/*CS* comparison are involved in many metabolic pathways, oxidoreductase activity, structural processes, and RNA processing (Fig. 3C-D; Supplementary Fig. 5). DEGs in SH conditions in *fru^LexA/4-40^* mutants include genes involved in supramolecular and defense processes (Fig. 3E-F; Supplementary Fig. 5). In comparison, SH *dsx^0683/1649^* mutants changed the expression of genes regulating cell-cell junctions and communication, development of the nervous system, and various signaling pathways (Fig. 3E-F; Supplementary Fig. 5). Overall, transcriptional profiles in both GH and SH brains revealed some genes to be exclusively impacted in either *fru^LexA/4-40^* or *dsx^0683/1649^* mutants, while others were affected in both mutants. These results suggest that distinct and shared transcriptional responses in the central brain are mediated by Fru^M^ and Dsx^M^.

Under the same housing condition, H3K4me3 or RNAPolII displayed differential distribution in *fru^LexA/4-40^* or *dsx^0683/1649^* mutants compared to *CS* in male brains (Fig. 3G-K; Fig. 6). Genes linked with H3K4me3 or RNAPolII DEPs were enriched in diverse GO terms (Fig. 3K). The *dsx^0683/1649^* mutants exhibited varying levels of H3K4me3 enrichment near genes related to DNA catalytic activity compared to *CS* under the GH condition, and showed differential enrichment of RNAPolII around genes involved in mating and reproductive behaviors, developmental and metabolic processes (Fig. 3K). While *fru^LexA/4-40^* mutants showed differential enrichment of RNAPolII at genes involved in synapse assembly or organization in the GH condition (Fig. 3K). *fru^LexA/4-40^* and *dsx^0683/1649^* mutants show both specific and shared influences on chromatin states in each housing condition compared to wild type (Fig. 3G-J). Consistent with their unique and shared roles in transcriptional regulation, these data continue to support distinct housing-specific changes in chromatin profiles around each gene mediated by Fru^M^ and Dsx^M^.

The comparison of DEGs and genes with DEPs in *fru^LexA/4-40^* and *dsx^0683/1649^* mutants relative to *CS* revealed concordant changes at both transcription and chromatin levels, specifically regarding H3K4me3 or RNAPolII enrichment. Notably, a significant number of genes with H3K4me3 DEPs in *dsx^0683/1649^* mutants corresponded to changes in transcription levels in the same direction, including many genes related to behaviors and DNA processes (Supplementary Fig. 3). Interestingly, *fru^LexA/4-40^* and *dsx^0683/1649^*mutants changed both the transcript level and chromatin level for circadian and sleep-related genes, *per* and *sand* (Supplementary Fig. 3; Fig. 7). Strikingly, the *sand* gene displayed dramatically decreased H4K3me3 enrichment and transcript level in *fru^LexA/4-40^* and *dsx^0683/1649^* mutants, with the RNAPolII binding level remaining the same (Supplementary Fig. 9). This might suggest either transcriptional priming or RNAPolII pausing at the *sand* gene locus in *fru^LexA/4-40^* and *dsx^0683/1649^* mutants, possibly due to the lack of H3K4me3 enrichment.

### Or47b circuit functions through Dsx^M^ in response to social isolation, whereas Or67d circuit functions through Fru^M^ in response to group housing

Our analyses have shown that *Or47b^2/3^*, *Or67d^GAL4/GAL4^*, *fru^LexA/4-40^*, and *dsx^0683/1649^* mutants all normalized the transcriptional differences between GH and SH comparison in unique ways (Fig. 2-4; Supplementary Fig. 7). Activation of pheromone-sensing neurons in the olfactory system activates ensembles of neurons within circuits driving social behaviors. This circuit activation can engage the functions of Fru^M^ and Dsx^M^ transcription factors expressed in a pheromone circuit-specific way to mediate the transcriptional and chromatin effects of social experience in the brain. To determine the neural and transcriptional cascades involved in this process and identify shared genes that are co-regulated by each pheromone circuit alongside Fru^M^ and Dsx^M^, we compared the DEGs and DEPs in each mutant (Fig. 4-6).

All mutants decreased the number of social experience-dependent DEGs in unique ways (Fig. 4A-D). In both *Or47b^2/3^* and *dsx^0683/1649^* mutants, the patterns of DEGs in response to social isolation are disrupted (Fig. 4D). This suggests that the Or47b circuits and Dsx^M^ are important for driving transcriptional responses to isolation. Notably, *dsx^0683/1649^* mutants show a more significant defect in social experience-induced transcriptional changes, both in terms of the number of genes affected and the level of transcriptional change observed (Fig. 4A-D). In contrast, in *Or67d^GAL4/GAL4^* and *fru^LexA/4-40^* mutants, transcriptional responses to group housing were largely disrupted (Fig. 4D), suggesting that Or67d circuits and Fru^M^ drive transcriptional responses to group housing. Again, *fru^LexA/4-40^* mutants exhibited a stronger defect in transcription. Notably, *Dsk*, which encodes a neuropeptide involved in regulating aggression and suppressing courtship behaviors (Agrawal et al., 2020; Wu et al., 2019), is the only gene statistically unaltered between grouped and isolated male brains across all wild-type and mutant genotypes (Fig. 4C; Supplementary Fig. 1D).

Comparing each mutant to *CS* under the GH condition revealed that *Or47b^2/3^* mutants showed the least number of DEGs among other mutants (Fig. 4E-F). In contrast, under the SH condition, *Or67d^GAL4/GAL4^* mutants showed the least number of DEGs compared to *CS* (Fig. 4I-J). With respect to wild-type brains, *fru^LexA/4-40^* mutants or *dsx^0683/1649^* mutants always had more dramatic effects on transcriptional profiles compared to Or mutants (Fig. 4E-L), supporting the hypothesis that *fru-* or *dsx-*positive central circuits likely integrate information from multiple pheromone circuits unexplored in this study.

We next aimed to determine whether we could identify functionally shared or segregated pathways connecting different pheromone circuits to the Fru^M^ and Dsx^M^ transcriptional functions in the central brain. We compared the DEGs in Or mutants to those in TF mutants versus *CS*. *Or47b^2/3^* mutants had a large number of DEGs that uniquely overlapped (268) with *dsx^0683/1649^* mutants under the SH condition (Fig. 4K). Meanwhile, a large number of DEGs in *Or67d^GAL4/GAL4^* and *fru^LexA/4-40^* mutant brains uniquely overlapped (197) in the GH condition (Fig. 4G). Among all the DEGs from comparisons of any mutant versus *CS*, a notable number showed altered expression in the same direction across all four mutants, or exhibited unique directional changes in specific mutants (Fig. 5). A significant number of the remaining DEGs showed interesting overlaps in expression patterns between Or (*Or47b* or *Or67d*) and TF (*fru^M^* or *dsx^M^*) mutants. We generally observed a trend indicating that, in both GH and SH conditions, the transcriptional effects on DEGs seen in *Or47b^2/3^* mutants are largely similar to those in *dsx^0683/1649^* mutants, and occasionally to *fru^LexA/4-40^* mutants (Fig. 5 gray frames). We saw opposing trends with *Or67d^GAL4/GAL4^* mutants, where the transcriptional effects generally aligned with *fru^LexA/4-40^*mutants, and occasionally with *dsx^0683/1649^* mutants (Fig. 5 gray frames).

**Figure 5:**
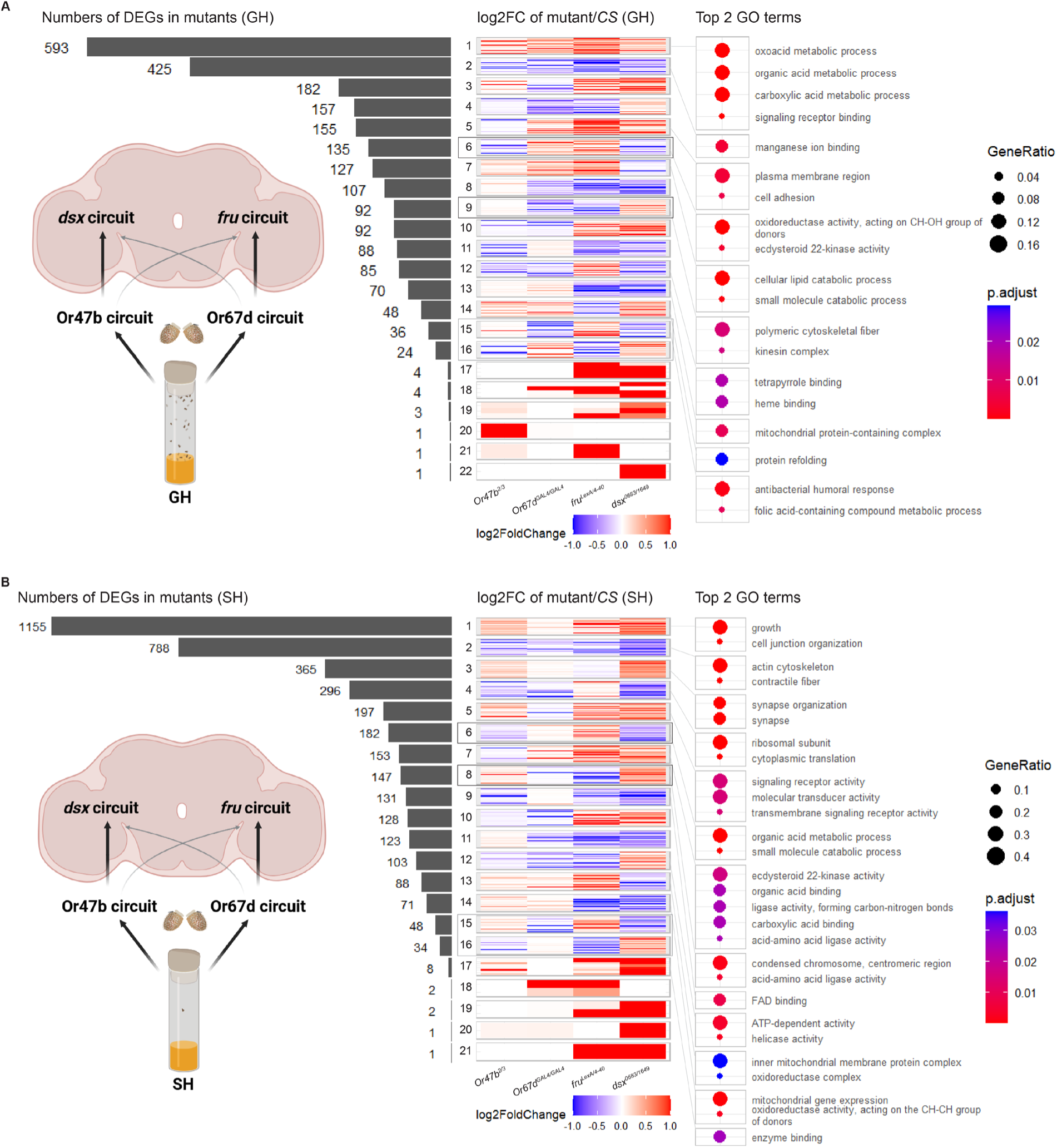
Or47b and Or67d circuits contribute to social modulation of gene expression through the function of Dsx^M^ and Fru^M^, respectively. **(A, B)** Bottom left: Models displaying pathways connecting Or47b or Or67d pheromone circuits to *fru* or *dsx* circuits in their transcriptional regulation on male fly brains under GH **(A)** or SH **(B)** condition. Left: Numbers of DEGs in mutant/*CS* comparisons across various combinations of log2FC in different directions. Middle: Heatmaps displaying log2FC of DEGs in mutant/*CS* comparisons across combinations of different directions. Dark gray frames indicate DEGs with log2FC in the same direction between *Or47b^2/3^*and *dsx^0683/1649^* mutants, as well as between *Or67d^GAL4/GAL4^* and *fru^LexA/4-40^*mutants. Light gray frames indicate DEGs with log2FC in the same direction between *Or47b^2/3^*and *fru^LexA/4-40^* mutants, and between *Or67d^GAL4/GAL4^* and *dsx^0683/1649^* mutants. Right: Top 2 GO terms for DEGs in mutant/*CS* comparisons across combinations of log2FC in different directions in the middle.

DEGs that changed in various combinations of directions were enriched in distinct GO terms (Fig. 5). Though DEGs showed enrichment for specific biological processes, molecular functions or cellular components in each mutant, under both housing conditions, DEGs in *Or47b^2/3^* and *dsx^0683/1649^* mutants shared many of the top ranked GO and KEGG terms (Supplementary Fig. 5). In contrast, DEGs in *Or67d^GAL4/GAL4^* and *fru^LexA/4-40^* mutants showed overlaps in GO and KEGG terms (Supplementary Fig. 5). These results highlighted potential functional connections between Or47b and Dsx^M^ neuronal circuits, particularly in isolated males, and between Or67d and Fru^M^ neuronal circuits especially in grouped males, suggesting that Or47b circuits might communicate with *dsx-* positive neurons in response to social isolation whereas Or67d circuits might function through *fru-*positive neural circuits in response to social enrichment.

At the chromatin level, genes with H3K4me3 or RNAPolII DEPs were mostly unique to each mutant, showing modest overlap with other genotypes (Fig. 6A-B). These genes were associated with distinct GO terms that were more genotype-specific (Fig. 6C). Figure 7 summarizes genes exhibiting concordant changes in both transcription and chromatin state (H3K4me3 or RNAPolII) in the same direction in each genotype. These findings suggest that transcriptional changes in these genes may be driven, at least in part, by broader chromatin remodeling events in the male brain induced by activation of different pheromone circuits in response to social experience, as well as the functions of Fru^M^ and Dsx^M^.

**Figure 6:**
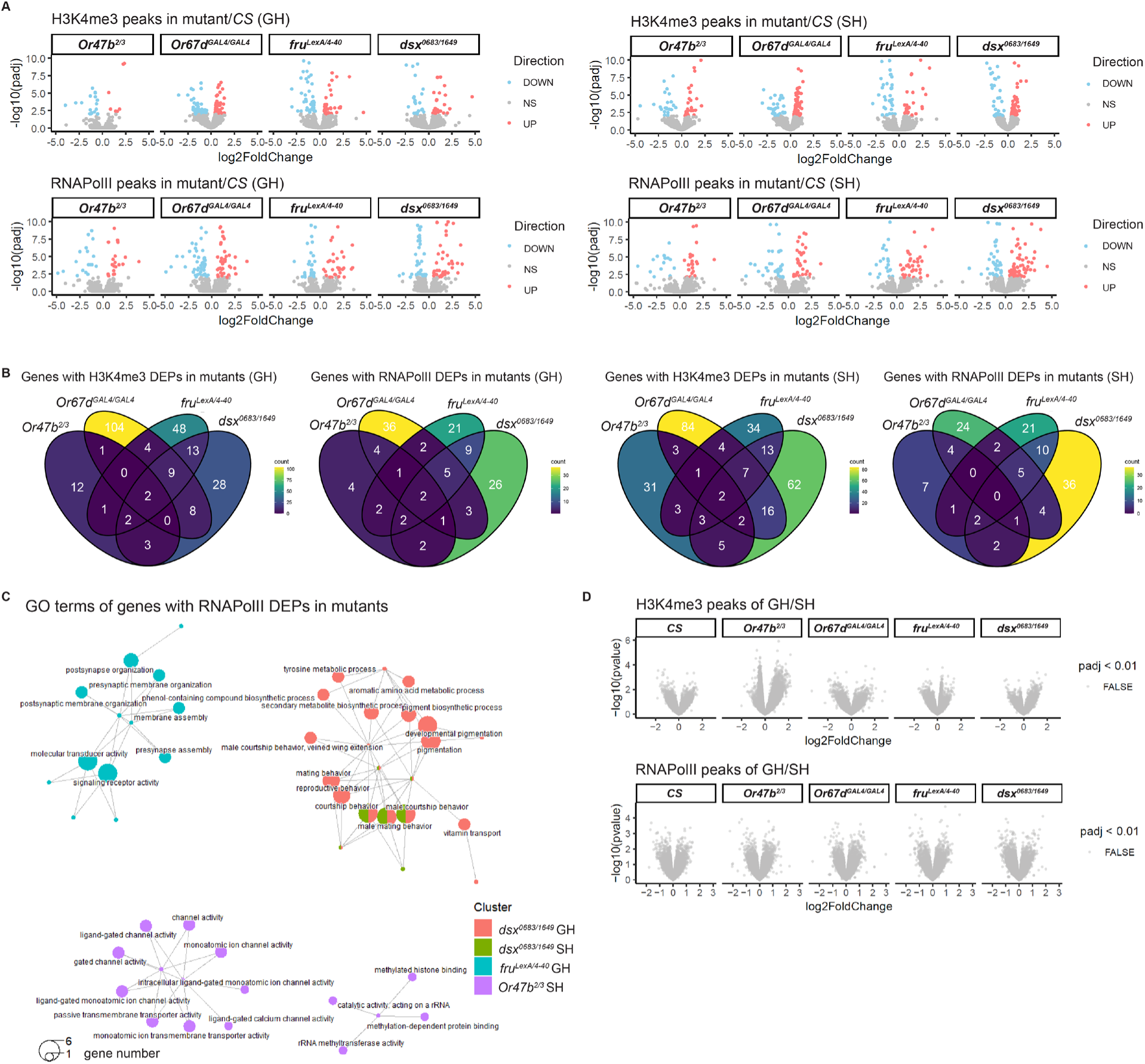
Comparisons of DEPs across Or and TF mutants in male fly brains in different social contexts. **(A)** Volcano plots showing log2FoldChange and –log10(padj) of mutant/*CS* comparisons for H3K4me3 or RNAPolII peaks under the same housing conditions. Peaks significantly enriched in mutants compared to *CS* are shown in red, while peaks significantly depleted are shown in blue (adjusted p-value < 0.01). **(B)** Venn diagram comparing genes with H3K4me3 or RNAPolII DEPs from mutant/*CS* comparisons under the same housing conditions. **(C)** Connection plots of GO terms for genes with RNAPolII DEPs from mutant/*CS* comparisons (q-value < 0.05). **(D)** Volcano plots showing log2FoldChange (GH/SH) and -log10(p-value) for H3K4me3 and RNAPolII peaks in each genotype. Gray labels peaks with adjusted p-value >= 0.01.

**Figure 7:**
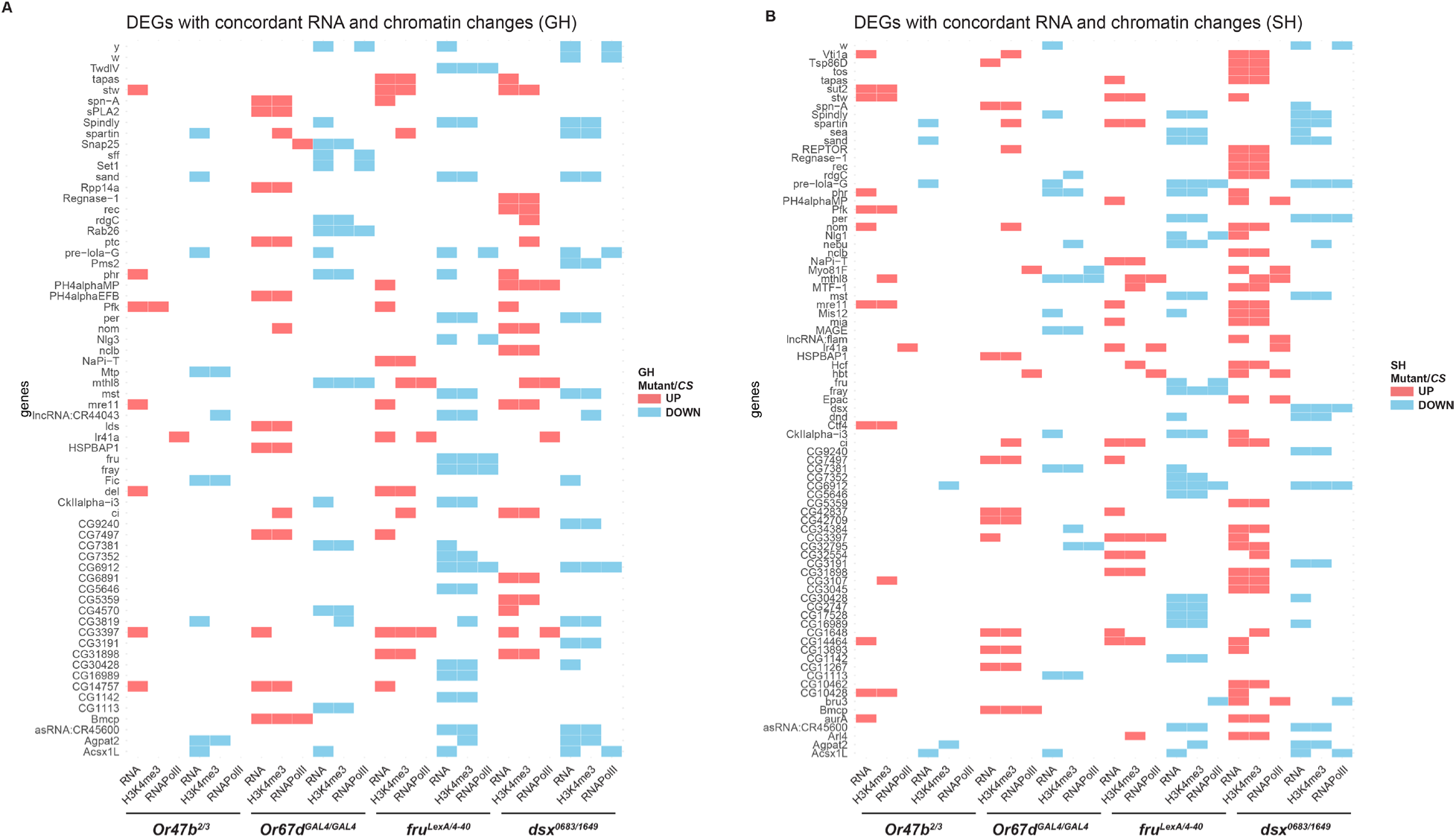
Concordant changes in transcription and chromatin in response to grouping or isolation. **(A, B)** Summary of genes exhibiting concordant changes in transcription and at least one active chromatin mark in each mutant/*CS* comparison under GH **(A)** or SH **(B)** conditions.

### Differential effects of pheromone circuits, Fru^M^, and Dsx^M^ on the social modulation of Fru^M^/Dsx^M^ target gene expression

Our results suggest that prolonged activation of pheromone circuits impacts the transcriptional activity of Fru^M^ or Dsx^M^-expressing neurons associated with sex-specific and/or social behaviors. This likely leads to changes in the expression of Fru^M^ and/or Dsx^M^ downstream target genes, contributing to the social modulation of behaviors. A comparison of Fru^M^ and Dsx^M^ binding sites in the genome revealed overlap among the genes regulated by either Fru^M^ or Dsx^M^ (Fig. 4D, H, L; Supplementary Fig. 6A-B) (Clough et al., 2014; Neville et al., 2014). Our analysis identified genes that are exclusively regulated/co-regulated by Fru^M^ and Dsx^M^ (Supplementary Fig. 6), which have various functions, including neural development and function, as well as modulation of circadian rhythms, metabolic processes, and signaling pathways (Clough et al., 2014; Neville et al., 2014).

We first analyzed the effect of housing on Fru^M^ and Dsx^M^ target DEGs, and how they changed in *fru^LexA/4-40^*mutants and *dsx^0683/1649^* mutants. We found that many of the Fru^M^/Dsx^M^ target genes were misexpressed primarily under one housing condition. Generally, *fru^LexA/4-40^* mutants exhibited defects in GH, while *dsx^0683/1649^* mutants showed defects in SH (Supplementary Fig. 6A). These DEGs contain binding sites for either Fru^M^ or Dsx^M^, or both, in their regulatory elements (Supplementary Fig. 6A). Under the same housing conditions, some of the target DEGs were either up-or down-regulated in the same direction in both mutants, albeit generally with one mutant having a more dominant effect (Supplementary Fig. 6B-D). While many co-regulated DEGs showed changes in the same direction, others were affected in opposite directions (Supplementary Fig. 6B-D). Some of the target DEGs were regulated exclusively by either Fru^M^ or Dsx^M^ (Supplementary Fig. 6B-D). These data suggest that social experience-induced changes in Fru^M^/Dsx^M^ target DEGs are mainly influenced by Fru^M^ in group-housed conditions, while Dsx^M^ primarily drives responses to social isolation. Furthermore, they reveal both similarities and differences in the mode of transcriptional regulation between Fru^M^ and Dsx^M^ on the expression of their target genes, where both can function as activators or repressors of transcription.

To map the neural pathways from the olfactory pheromone-sensing neurons to the regulation of Fru^M^, Dsx^M^ and their target DEGs in the central brain, we compared the log2 fold change of these DEGs in *Or47b^2/3^*and *Or67d^GAL4/GAL4^* mutants as well. The effects of *Or47b^2/3^* and *Or67d^GAL4/GAL4^* mutants on the Fru^M^/Dsx^M^ target DEGs were not as dramatic as *fru^LexA/4-40^* or *dsx^0683/1649^* mutants (Supplementary Fig. 6B-D). We identified some DEGs that were either altered in the same direction in all 4 mutants, or in the opposite direction only in a single mutant (Supplementary Fig. 6C-D). Similar to prior results, in both GH and SH conditions, *Or47b^2/3^* mutants show transcriptional effects on target DEGs similar to *dsx^0683/1649^* mutants, and occasionally with *fru^LexA/4-40^* mutants (Supplementary Fig. 6C-D). In contrast, *Or67d^GAL4/GAL4^* mutants generally align with *fru^LexA/4-40^* mutants, and occasionally *dsx^0683/1649^* mutants (Supplementary Fig. 6C-D). These results are in agreement with the results above that suggest Or47b and Or67d circuits predominantly work through Fru^M^ and Dsx^M^, respectively.

In an accompanying study, we discovered that social experience influences the expression of circadian-related genes (i.e., *cry, Clk, Pdf,* and *Sik3*) in the central brain (Du et al., 2025). Interestingly, many of these circadian genes are Fru^M^/Dsx^M^ target genes with Fru^M^ and/or Dsx^M^ binding in their upstream regulatory elements (Clough et al., 2014; Neville et al., 2014). Due to the pleiotropic effects of the circadian system on behavior and physiology, social experience-dependent alterations in Fru^M^/Dsx^M^ transcriptional cascades and the expression of downstream target circadian genes may drive the impact of social experiences on behaviors. Indeed, knockdown of *fru^M^* and mutants in various circadian genes either amplified or reduced the effects of group housing-driven changes in courtship (Clough et al., 2014; Neville et al., 2014). These results indicate that changes in circadian gene expression, influenced by social experience, may arise as a consequence of alterations in the levels or functions of Fru^M^ or Dsx^M^. To test this hypothesis and more strictly temporally control transcriptional changes in circadian genes, we sampled RNA at ZT7-8 from grouped and isolated male heads in different mutants and performed qRT-PCR for *Clk, tim,* and *cry* with control genes (*TSG101*, *RpS29*, *Hmt4-20*, and *Dsk*). We found that expression of *Clk* and *cry* trended towards an increase in response to isolation (Supplementary Fig. 8A) (Du et al., 2025). Under both social conditions, *tim* and *cry* showed a decrease in *dsx^0683/1649^* mutants (Supplementary Fig. 8B). *Or67d^GAL4/GAL4^* mutants showed the strongest decrease in the expression of *cry*, in the same direction as observed in *dsx^0683/1649^* mutants (Supplementary Fig. 8B). *dsx^0683/1649^* mutants also showed a slight increase in *Clk* expression in GH brains, which eliminated the difference in *Clk* expression observed between GH and SH *CS* brains (Supplementary Fig. 8A-B). In contrast, in *fru^LexA/4-40^* mutants, *cry* expression decreased in SH brains, while *tim* expression increased in GH brains (Supplementary Fig. 8B). These results suggest that Fru^M^ and Dsx^M^ differentially regulate their target genes, i.e., circadian genes, in the head, and social experience can modulate the expression of the circadian genes in response to the differential activation of different pheromone circuits and transcriptional function of Fru^M^ and Dsx^M^.

### Differential effects of pheromone circuits, Fru^M^, and Dsx^M^ on social modulation of behaviors

Considering the transcriptional changes in the brains of mutants of *Or67d*, *Or47b*, *fru^M^,* and *dsx^M^*, we wanted to explore their effects on the social modulation of behaviors. Disrupting the function of Or47b or Or67d affects the detection of diverse social cues, while loss of Fru^M^ and Dsx^M^ function disrupts the processing of social signals in the central social circuits (Dauwalder, 2011; Dweck et al., 2015; Ha and Smith, 2006; Kurtovic et al., 2007; Lin et al., 2016; Vrontou et al., 2006; Yamamoto and Koganezawa, 2013). Previous studies have shown that social experience can reset the circadian clock in *Drosophila*, and grouped *CS* males show increased sleep and decreased locomotion compared to isolated *CS* males (Levine et al., 2002; Li et al., 2021; Zhao et al., 2024). Therefore, we assessed sleep and locomotion activity in *Or47b^2/3^*, *Or67d^GAL4/GAL4^*, *fru^LexA/4-40^*, and *dsx^0683/1649^* mutants to determine their contribution to social experience-dependent changes in behaviors (Fig. 8; Supplementary Fig. 10). We confirmed that GH *CS* males increased sleep and decreased locomotion compared to SH *CS* males (Fig. 8A-D). Both *fru^LexA/4-40^*and *dsx^0683/1649^* mutants diminished the differences in total sleep and locomotion between GH and SH males (Fig. 8A-D), indicating that they both play a role in the social modulation of these behaviors. In *dsx^0683/1649^* mutants, such elimination resulted from both a decreased GH sleep level and an increased SH level (Supplementary Fig. 10A-D). In *fru^LexA/4-40^* mutants, the sleep pattern showed a general decline in GH (Supplementary Fig. 10A-D). We also noted further differences in the transition of sleep from nighttime to daytime in *fru^LexA/4-40^* mutants (Fig. 8A). In particular, the activity bout at the transition from nighttime to daytime was not disrupted in *fru^LexA/4-40^* mutants as in other genotypes (Fig. 8A). This also agrees with their locomotion performance, in which *fru^LexA/4-40^* mutants showed an overall decrease in activity peaks in the transition, and a less significant difference in locomotion between SH and GH flies (Fig. 8C). *Or47b^2/3^* and *Or67d^GAL4/GAL4^* mutants did not dramatically alter sleep patterns or trends between GH and SH (Fig. 8A-B). However, *Or47b^2/3^* mutants decreased locomotion in SH flies (Fig. 8C-D; Supplementary Fig. 10E-H). This result aligns with transcriptional disruptions in genes involved in locomotion of *Or47b^2/3^* mutants in SH brains (Fig. 2B). These findings suggest that Fru^M^ and Dsx^M^ play essential but distinct roles in the social modulation of sleep and locomotion behaviors in isolated and grouped flies, with a potential role for Or47b in mediating changes in locomotion in SH flies.

**Figure 8:**
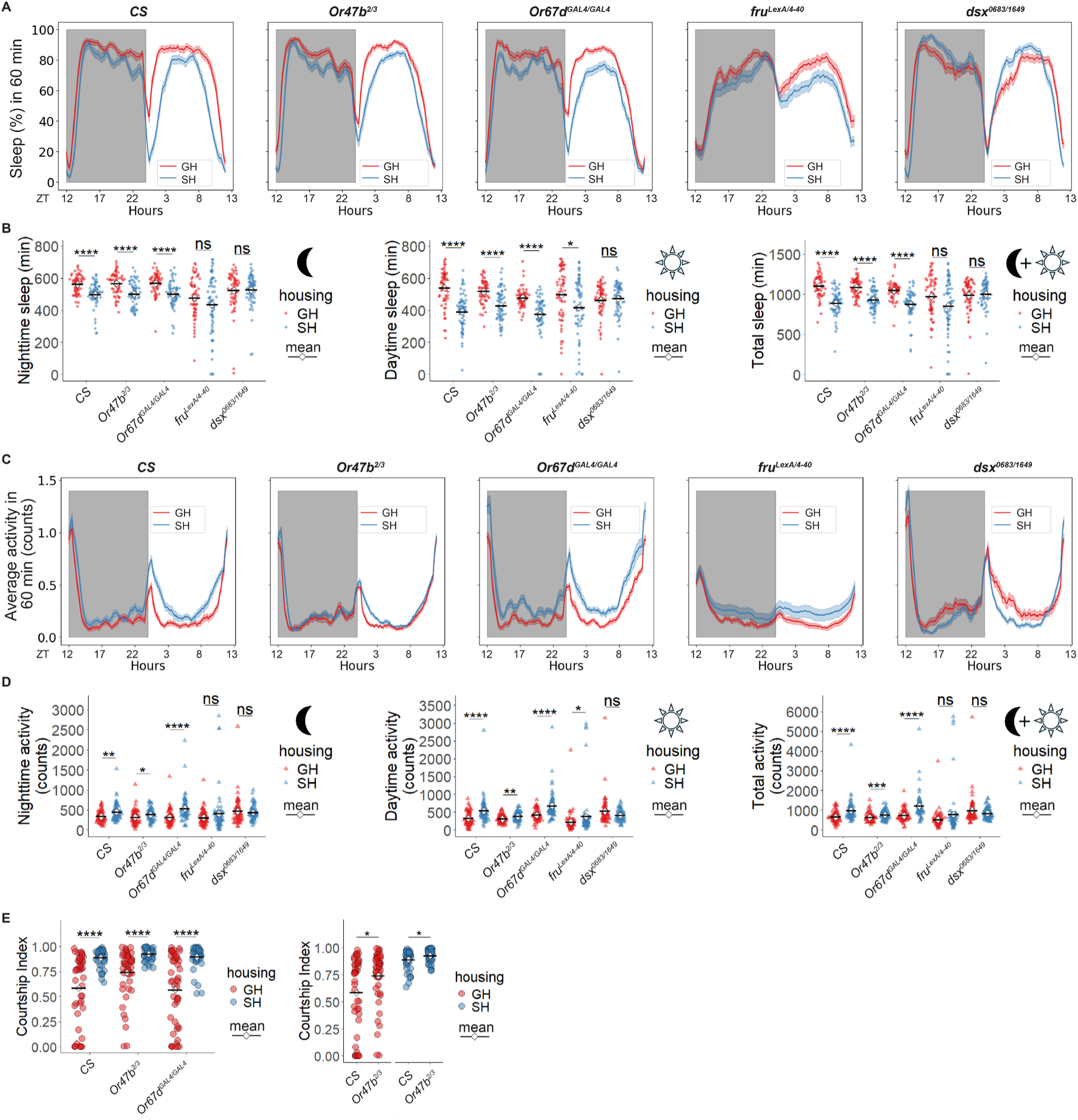
Or47b and Or67d pheromone circuits, Fru^M^, and Dsx^M^ have differential effects on social modulation of behaviors. **(A)** Sleep profiles of male flies from each genotype raised under GH or SH conditions, shown as the average proportion of sleep time per 60-minute interval during a 24-hour 12:12 dark-light cycle. Mean ±SEM. **(B)** Quantification of nighttime sleep, daytime sleep, and daily total sleep in **(A)**. Statistical comparisons were performed using the Wilcoxon test. P-values were adjusted for multiple comparisons across genotypes using the Benjamini–Hochberg method. **(C)** Locomotion activity profiles of male flies from each genotype raised under GH or SH conditions, shown as the average counts of beam crossings per 60-minute interval during a 24-hour 12:12 dark-light cycle. Mean ±SEM. **(D)** Quantification of nighttime activity, daytime activity and daily total activity in **(C)**. Statistical comparisons were performed using the Wilcoxon test. P-values were adjusted for multiple comparisons across genotypes using the Benjamini–Hochberg method. *P_adj<0.05, **P_adj<0.01, ***P_adj<0.001, ****P_adj<0.0001. (E) Male-female single pair courtship assay using *CS*, *Or47b^2/3^*, and *Or67d^GAL4/GAL4^* males raised under GH and SH conditions. Statistical comparisons were performed using the Wilcoxon test between GH and SH for each genotype (left) and the comparison between *Or47b^2/3^* and *CS* under the same housing conditions (right). P-values were adjusted for multiple comparisons across genotypes (left) or housing conditions (right) using the Benjamini–Hochberg method. Significance of adjusted P-value (P_adj): *P_adj < 0.05, **P_adj < 0.01, ***P_adj < 0.001, ****P_adj < 0.0001.

Since Or47b, Or67d, Fru^M^, and Dsx^M^ circuits are all related to courtship behaviors, we also performed single-pair male-female courtship assays to determine their effect on social modulation of courtship. Previous studies and our accompanying study showed that *fru^M^*and *dsx^M^* are necessary for normal courtship behaviors in male flies, and *fru^M^* expression in the central brain neurons is necessary for mediating the social experience-induced changes in male courtship behaviors (Du et al., 2025). The role of *fru^M^* regulation with social experience is diverse: in some neurons, altered Fru^M^ function enhances courtship behavior in SH males, while in other neurons it decreases courtship in response to GH (Du et al., 2025). Since social experience can influence gene regulation and behavior, and given that mutations in *Or47b* and *Or67d* affect the transcriptional profile and chromatin state of wild-type brains, we investigated whether either the Or47b or Or67d pheromone circuits play a role in how social experience modulates courtship behaviors. We quantified male courtship behaviors in *CS*, *Or47b^2/3^* and *Or67d^GAL4/GAL4^* mutants (Fig. 8E). Loss of *Or47b* function increased courtship in GH mutant males, though it did not completely eliminate the difference between GH and SH (Fig. 8E). The *Or67d^GAL4/GAL4^* mutants did not alter the courtship vigor in either social context (Fig. 8E). These results suggest that Or47b circuits contribute to courtship suppression in grouped males. Overall, our results suggest unique contributions of olfactory pheromone pathways and transcription factors critical for integrating social information on transcriptional and chromatin profiles are accompanied by equally unique effects on social modulation of various behaviors.

## Discussion

A substantial body of evidence demonstrates that social environments, whether isolation or enrichment, have lasting consequences for cognition, physiology, and behavior (Cacioppo and Hawkley, 2009; Dow-Edwards et al., 2019; Jeste et al., 2020). Loneliness, in particular, has been consistently linked to heightened risk and severity of neuropsychiatric and neurodegenerative disorders, establishing it as a pressing public health concern (General, 2023; Jeste et al., 2020; Lin, 2023). Nevertheless, while the behavioral outcomes of different social experiences are well recognized, the molecular pathways and neural circuits mediating these effects remain insufficiently understood. In this study, we investigated how transcription patterns and chromatin marks in the brains of male *Drosophila* change in response to social experience. To determine the neural and transcriptional pathways responding to social experience, we examined mutants in two pheromone receptors, Or47b and Or67d, along with two transcription factors, Fru^M^ and Dsx^M^, that influence sex-specific and social behaviors. The results show that *Or47b^2/3^*, *Or67d^GAL4/GAL4^, fru^LexA/4-40^*, and *dsx^0683/1649^* mutants significantly reduced the transcriptional responses to social experience, suggesting that social modulation of gene expression involves all four genes and their associated circuits. *Or47b^2/3^* and *Or67d^GAL4/GAL4^* mutants exhibited opposing effects on DEGs that are altered by social experience in wild type. Similar opposing functional effects were also observed in *fru^LexA/4-40^* and *dsx^0683/1649^* mutants. We found functionally segregated transcriptional cascades from individual pheromone circuits that engaged the function of either Fru^M^ or Dsx^M^ in the central brain. Mainly, we found that Or47b circuits generally function through Dsx^M^ in response to isolation, while Or67d circuits work in group-housed males through Fru^M^ to mediate social experience-dependent modulation of transcriptional responses. The RNA profiles from the brains of *fru^LexA/4-40^* and *dsx^0683/1649^*mutants also helped us identify both activation and repression functions for Fru^M^ and Dsx^M^, in addition to their shared and exclusive functions in mediating their effects on gene expression and behaviors in different housing conditions. And finally, we found that social experience-dependent changes in sleep and locomotion are generally eliminated in *fru^LexA/4-40^* and *dsx^0683/1649^* mutants, with moderate effects of pheromone receptors on some of the behaviors under certain social conditions. Given that many of the DEGs are direct targets of Fru^M^ and/or Dsx^M^, our study reveals how changes in social and pheromone circuit activity with social experience can alter transcriptional cascades reprogramming brains and behaviors.

One surprising finding from our study is that, although we identified DEGs between grouped and isolated male brains of wild-type flies, we did not observe dramatic changes in chromatin patterns of H3K4me3 and RNAPolII across any of the genotypes under different housing conditions. However, we detected numerous changes to the enrichment of H3K4me3 and RNAPolII in each mutant compared with wild-type brains under the same housing conditions. Some of these changes in chromatin were accompanied by concordant changes at the transcription level. Combinatorial function of multiple histone modifications together with RNAPolII is thought to be intimately connected to transcriptional states (Goldberg et al., 2007; Jenuwein and Allis, 2001; Schones and Zhao, 2008). For instance, H3K4me3 and H3K27ac are considered marks for actively transcribed genes, while H3K27me3 is enriched around transcriptionally repressed genes (Boros, 2012; Lawrence et al., 2016; Schones and Zhao, 2008; Zhang et al., 2021; Zhou et al., 2011). However, many studies have also shown discrepancies between the enrichment of specific chromatin marks and transcription responses (Agrawal et al., 2019; Brovkina et al., 2021; Kiani et al., 2022). For example, circadian analysis of transcriptional and chromatin states in mammalian cells throughout the day revealed that such correlations might represent an oversimplified view (Koike et al., 2012). This study found that while both transcription and chromatin states displayed circadian patterns, numerous genes exhibited changes in chromatin patterns without affecting transcription levels and vice versa. Comparison of transcriptional changes and histone marks in dopaminergic neurons in flies raised in grouped versus isolated conditions also revealed only a small correlation in a limited number of genes (Agrawal et al., 2019). They also discovered that many highly expressed genes showed unexpected increases in the enrichment of the repressive H3K27me3 mark, alongside decreases in active chromatin marks (Agrawal et al., 2019). Similarly, chromatin accessibility and gene expression in *fru*+ versus *fru*- cells sorted from male and female fly brains show little overlap (Brovkina et al., 2021).

There are several explanations for the discrepancy we and others observe in transcript levels and chromatin mark enrichment. One possibility is cellular heterogeneity. Changes in chromatin states may occur in a small subset of cells in response to social experience. However, when analyzing the whole brain, these subtle changes can be obscured by other cells where such changes are not happening. Single-cell multiome ATAC and gene expression profiling techniques can simultaneously determine gene expression and chromatin landscape from the same cell, potentially increasing the resolution and confidence of molecular analysis (Yamagishi et al., 2024). An alternative explanation considers the transcriptional and chromatin states of a gene as it transitions through active, repressed, and poised states. Each of these states may be triggered by complex combinations of chromatin marks that accumulate around genes as cells respond to various signals and pathways. Transitioning between active and repressed transcriptional states may involve intermediate states as chromatin modifications build up around responsive genes. These conditions could lead to intermediate states where transcription decreases alongside an increase in repressive marks, while active marks remain unchanged. Conversely, transcription might increase due to the loss of a repressive mark, rather than the gain of an active mark. In this study, we only investigated two active marks, yet marks for repressed/closed chromatin are also crucial for regulating gene expression. Studies on mammalian cells demonstrated that the promoters of some transcriptionally repressed genes were marked with both active (H3K4me3) and repressive (H3K27me3) epigenetic marks (Mikkelsen et al., 2007). These and numerous other studies indicate that chromatin states, as signaled by histone marks or accessibility, only offer a probabilistic prediction of transcription (Corbett, 2018; Fu et al., 2025). Changes in gene expression result from the combined regulation of various factors, including chromatin accessibility, histone modifications, transcription factors, and post-transcriptional modifications (Corbett, 2018; Fu et al., 2025). Comprehensive studies that analyze gene expression across all features will lead to a better understanding.

Despite the discrepancy in chromatin and transcriptional states, we did identify a small list of genes with concordant changes in transcript levels and active chromatin mark enrichment, like the *sand* gene. The *sand* gene, which encodes a KCNK18/TRESK homolog potassium channel, plays a crucial role in regulating various aspects of *Drosophila* behavior and physiology. It is involved in sleep regulation within the dorsal fan-shaped bodies (Pimentel et al., 2016) and cardiac function (Abraham et al., 2018). Additionally, it has been shown to operate in cortical glial cells to help control stress-induced seizures (Weiss et al., 2019). Considering the important role of the *sand* gene and the notable alterations in its transcriptional and chromatin state in *fru^LexA/4-40^* and *dsx^0683/1649^* mutants, it is possible that Sand may play a role in mediating some of the impaired neural, behavioral, and physiological processes observed in these mutants. Further studies are necessary to explore this hypothesis.

In a related paper, we demonstrated that the expression of *fru^M^*and core circadian genes, which are potentially regulated by Fru^M^ and Dsx^M^, is altered by social experience and essential for changes in courtship behaviors (Du et al., 2025). These results suggest that social experience might exert its effect on behavioral modulation through changes in circadian state. In this study, we further strengthen these findings by highlighting the roles of both Fru^M^ and Dsx^M^ in mediating the effects of social experience on additional behaviors known to be under circadian control, such as sleep and locomotion. We discovered that the Or47b circuits influence the social modulation of locomotion and courtship behaviors in isolated and grouped flies, respectively, correlating with changes in genes that mediate these behaviors in *Or47b^2/3^* mutant brains. Importantly, we identified the different effects of Or47b and Or67d pheromone circuits and how they interact with the functions of Fru^M^ and Dsx^M^ to regulate their target genes under various social conditions. Given that the regulation of circadian gene networks is complex and participates in the modulation of various behaviors through changes in arousal states, understanding the diverse impacts of Fru^M^ and Dsx^M^ on social modulation of target circadian gene expression in the brain is both important and challenging. Fru^M^, Dsx^M^, and various circadian genes are expressed in different neuronal and non-neuronal (glial and fat body) cell populations in the brain in different combinations. The cellular location of social experience-dependent changes in Fru^M^ and Dsx^M^ function and target gene expression remains unknown. Further studies are essential to identify the specific cells in which Fru^M^ and Dsx^M^ mediate transcriptional differences in circadian genes in response to social experience. We anticipate that these studies will provide a clearer understanding of how social experience modifies the complex gene regulatory networks for circadian genes downstream of Fru^M^ and Dsx^M^ and how these modifications alter behaviors.

Overall, our study uncovers how social experience alters transcription and chromatin states in the brain, and provides insights into the roles played by pheromone circuits and transcription factors that influence molecular and behavioral responses to social experience. In addition to modulating behavioral responses, social experience and circadian pathways also influence many physiological responses. These are clearly represented in our transcriptome dataset as social experience-dependent DEGs implicated in immune responses, metabolism, signal transduction, homeostasis, and developmental pathways. The activity of pheromone circuits, along with the functions of Fru^M^ and Dsx^M^, may also play a role in how social experience influences physiological responses through changes in downstream target genes. Future studies will reveal how social experience impacts additional behavioral and physiological responses, and the effects of different pheromone circuits (from both olfactory and gustatory systems), as well as transcriptional and circadian pathways, on these responses.

## Methods

### Drosophila strains and husbandry

Flies were raised on standard fly food from Archon Scientific Company at room temperature. Male flies used in all experiments were 7-day-old. For male-female single-pair courtship behavior assays, unmated males were used as the test fly, and 6-day-old *w^1118^* virgin females were used as the target fly. Specific genotypes are listed here (Tweedie et al., 2009): wild-type *CS* (*Canton-S*), *Or47b* mutant (*w^+^/Y; Or47b^2/3^;+/+*) (Wang et al., 2011), *Or67d* mutant (*w^+^/Y; +/+; Or67d^GAL4/GAL4^*) (Kurtovic et al., 2007), *fru* mutant (*w^+^/Y; +/+; fru^LexA/4–40^*) (Anand et al., 2001; Mellert et al., 2010), and *dsx* mutant (*w^-^/Y; +/+; dsx^f00683-d07058/f01649-d09625^*) (Chatterjee et al., 2011).

### Social isolation and group housing setup

Dark pupae (80–100 hours old) were picked out and separated by sex. For the social isolation (single housing, SH) condition, each pupa was placed into individual vials, allowed to eclose alone, and aged to 7 days. For the group housing (GH) condition, more than 30 pupae of the same sex were collected and placed into food vials, and 30 newly eclosed and unmated flies with correct genotypes were collected and put in a fresh new vial the next day of the collection day (day 0). All *w^1118^* virgin females used for the courtship assay were raised under GH conditions.

### Bulk tissue RNA sequencing

#### Brain dissection and RNA extraction

Fly brains were dissected in cold RNase-free 1X PBS and transferred immediately to RNAlater (Sigma-Aldrich, R0901) on ice. Each genotype under the GH or SH conditions has three biological replicates, and each biological replicate contains 30 male fly brains. After the dissection, RNAlater with the sample was diluted 1:2 with 1X RNase-free PBS and centrifuged for 1 min at max speed, and the supernatant was removed. Brain samples were washed with 1 mL 1X RNase-free PBS and centrifuged for 1 min at max speed. The supernatant was removed and replaced with 300 μL of TRIzol reagent (Invitrogen, 15596026). To extract RNA, samples were ground sufficiently and processed with QIAGEN Shredder kit (QIAGEN, 79654), and RNeasy Mini Kit (QIAGEN, 74104).

DNase I (TURBO DNA-free Kit, Invitrogen, Thermo Fisher Scientific AM1907) was applied to remove genomic DNA after the RNA extraction.

#### Library preparation and sequencing

Around 500 ng total RNA in each sample was prepared for library preparation. KAPA Stranded mRNA-Seq Kit was used to generate mRNA libraries. Libraries were sequenced on the NextSeq 2000 sequencer, generating 75-bp or 100-bp paired-end reads.

#### Bulk RNA sequencing data analysis

Raw fastq files were processed with the standard nf-core/rnaseq pipeline (Ewels et al., 2020). Briefly, reads were mapped to the *Drosophila melanogaster* BDGP6.46 genome with STAR, and read counts were generated using the Salmon method. The following analysis was performed in R (R Core Team, 2013; Wickham and Sievert, 2009). DEG analysis was performed with DESeq2 (Love et al., 2014). All control and mutant samples were incorporated into a single DESeqDataSet (dds) object for differential expression analysis. The Wald test was applied to the comparison between GH and SH of the same genotype, and the comparison between mutant and *CS* under the same housing conditions. Gene ontology (GO), Reactome, and KEGG analyses were performed using the clusterProfiler library (Yu et al., 2012).

### CUT&RUN

#### Brain dissociation and chromatin immunoprecipitation

Fly brains were dissected in dissection buffer (Schneider′s Insect Medium, Sigma-Aldrich S0146 with 1% BSA) and transferred into cold dissection buffer in 1.5 ml tubes on ice. Each genotype under the GH or SH conditions has three biological replicates. For H3K4me3 samples, fifteen male brains were used per sample, and for RNAPolII samples, thirty male brains were used per sample. After completing the dissection, the old dissection buffer was replaced with a fresh one. Collagenase Type I (Gibco, 17018029) stock solution was added directly into samples for a final 2mg/mL concentration. Samples were incubated at 37 °C for 20 minutes. During the incubation, samples were dissociated by gentle pipetting 20-30 times every 7 min with P200. The collagenase solution was removed after centrifugation at 300 xg for 5 min. Cells were washed with wash buffer supplied in CUTANA™ ChIC/CUT&RUN Kit (EpiCypher, 14-1048), and were centrifuged at 300 xg for 5 min. The old wash buffer was replaced with fresh wash buffer, and the cells were resuspended and processed according to the instructions in the user manual. 1 ul of antibody (anti-H3K4me3 antibodies supplied by the CUT&RUN Kit, and anti-RNAPolII antibody from MilliporeSigma 05-623) was added per sample.

#### Library preparation and sequencing

CUTANA™ CUT&RUN Library Prep Kit (EpiCypher, 14-1001, 14-1002) was used to prepare libraries. The quality of libraries was examined by the D5000 ScreenTape station. Libraries were sequenced on the NextSeq 2000 sequencer, generating 75-bp paired-end reads.

#### CUT&RUN data analysis

Raw fastq files were processed using the standard nf-core/cutandrun pipeline (Ewels et al., 2020). Briefly, reads were trimmed by Trim Galore and aligned to the *Drosophila melanogaster* dmel-all-chromosome-r6.45 genome with Bowtie 2. Peak calling was performed by MACS2. The following analysis was performed in R (R Core Team, 2013; Wickham and Sievert, 2009). Counts of peaks were generated by csaw::regionCounts (Lun and Smyth, 2014, 2015). DEP analysis was performed with DESeq2 (Love et al., 2014). All control and mutant samples were incorporated into a single DESeqDataSet (dds) object for DEP analysis. Wald test was applied to the comparison between GH and SH of the same genotype, and the comparison between mutant and *CS* under the same housing conditions. ChIPseeker was used to annotate DNA peaks with genes. Gene ontology (GO), Reactome, and KEGG analyses were performed using the clusterProfiler library (Yu et al., 2012).

### Quantitative reverse transcription PCR (RT-qPCR)

The RT-qPCR protocol was modified based on the previous protocol (Li et al., 2016). For each housing condition, three to four biological replicates were prepared separately, with each replicate containing 15-20 heads from male flies. Heads were dissected at ZT7-ZT8 on fly pads and immediately transferred into 300 ul TRIzol (Invitrogen, 15596026) on ice. RNA extraction and DNase treatment were as described above. cDNA was synthesized through reverse transcription reaction using approximately 200 ng of total RNA with the SuperScript IV First-Strand Synthesis Kit (Invitrogen, 18091050) and poly d(T) primers. qPCR was performed using the FastStart Essential DNA Green Master kit (Roche, 06924204001) on LightCycler® 96 instrument (Roche, 05815916001). Primers used are listed in Table 1. The expression level was calculated by the ΔCt method using *TSG101* as the standard gene. The calculation was performed in R. For each genotype, relative expression between GH and SH male flies was compared using two-sided independent t-tests. For each housing condition, differences among genotypes were assessed using one-way ANOVA followed by Tukey’s HSD post-hoc test for pairwise comparisons. In both analyses, raw p-values were adjusted for multiple testing across all comparisons using the Benjamini–Hochberg procedure to control the false discovery rate. Adjusted p-values (*p.adj*) were used to determine statistical significance.

**Table 1.**
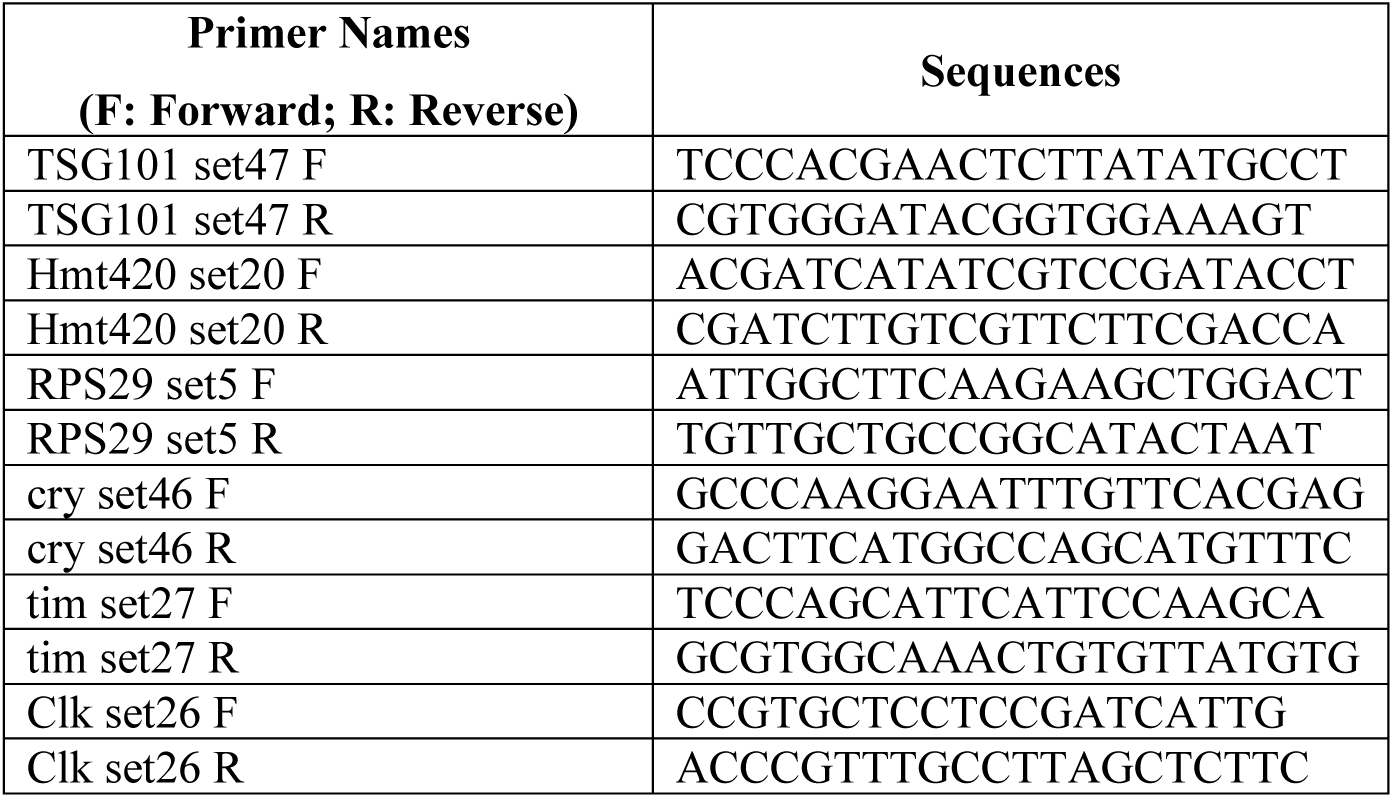
Primers used in RT-qPCR experiments.

### Male-female single-pair courtship behavior assay setup and analysis

Courtship behavior assays were performed during ZT 7 – ZT 10 as described (Du et al., 2025). Briefly, one unmated male and one virgin *w^1118^* female were gently aspirated to each arena of an unused agarose chamber, separated by barriers before the start of the recording. When all flies were loaded, the chamber was placed under a camera with a stable, consistent light source. Barriers were removed to allow the two flies to meet each other, and video recording was started immediately. The trajectory of each fly was tracked by FlyTracker (Eyjolfsdottir et al., 2014) on MATLAB. Fly identity and the start time of copulation were determined manually. JAABA was trained and used to automatically annotate other courtship behaviors (Kabra et al., 2013). For each courtship behavior video, only the first 10 minutes were used for the scoring of the courtship index (CI) (copulation time included):

CI = (the time a male fly spends on courtship)/(10 min)

### Sleep and locomotion behavior assay setup and analysis

Flies were grouped or isolated for 7 days, and then loaded individually into glass tubes containing food. The activity of each fly was monitored using the Drosophila Activity Monitor (DAM) system (TriKinetics, Waltham, MA) at 25 °C under 12-h light-dark (LD) cycles. The DAM system records activity counts *Ct*(*t*) (the number of counts each fly passes the beam) every 30 seconds for two days after the loading to capture one full LD cycle.

Sleep is defined as 5 consecutive minutes or longer with no beam passing.

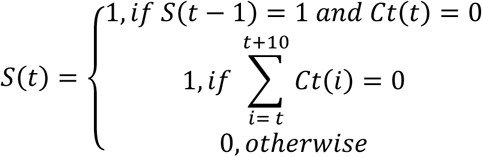

Sleep traces (%) were generated by averaging the sleep data over a duration of 60 minutes.

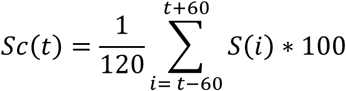

Locomotion traces (counts) were generated by averaging the activity counts over a duration of 60 minutes.

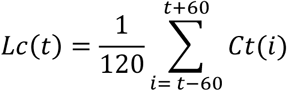

## Acknowledgements

We would like to thank members of the Volkan lab for discussions and help with the manuscript. We thank Dr. E. Josie Clowney (University of Michigan, Ann Arbor) for kindly sharing their CUT&RUN protocols. We thank Bloomington Stock Center and Flybase for their services.

## Author contributions

Conceptualization: CD and PCV. Investigation: CD, LJ, and SS with the help from SO and SR. Analysis/interpretation of data: CD, JESF, LJ, CDJ, and PCV. Manuscript—original draft: CD and PCV.

## Figures

**Supplementary Figure 1:**
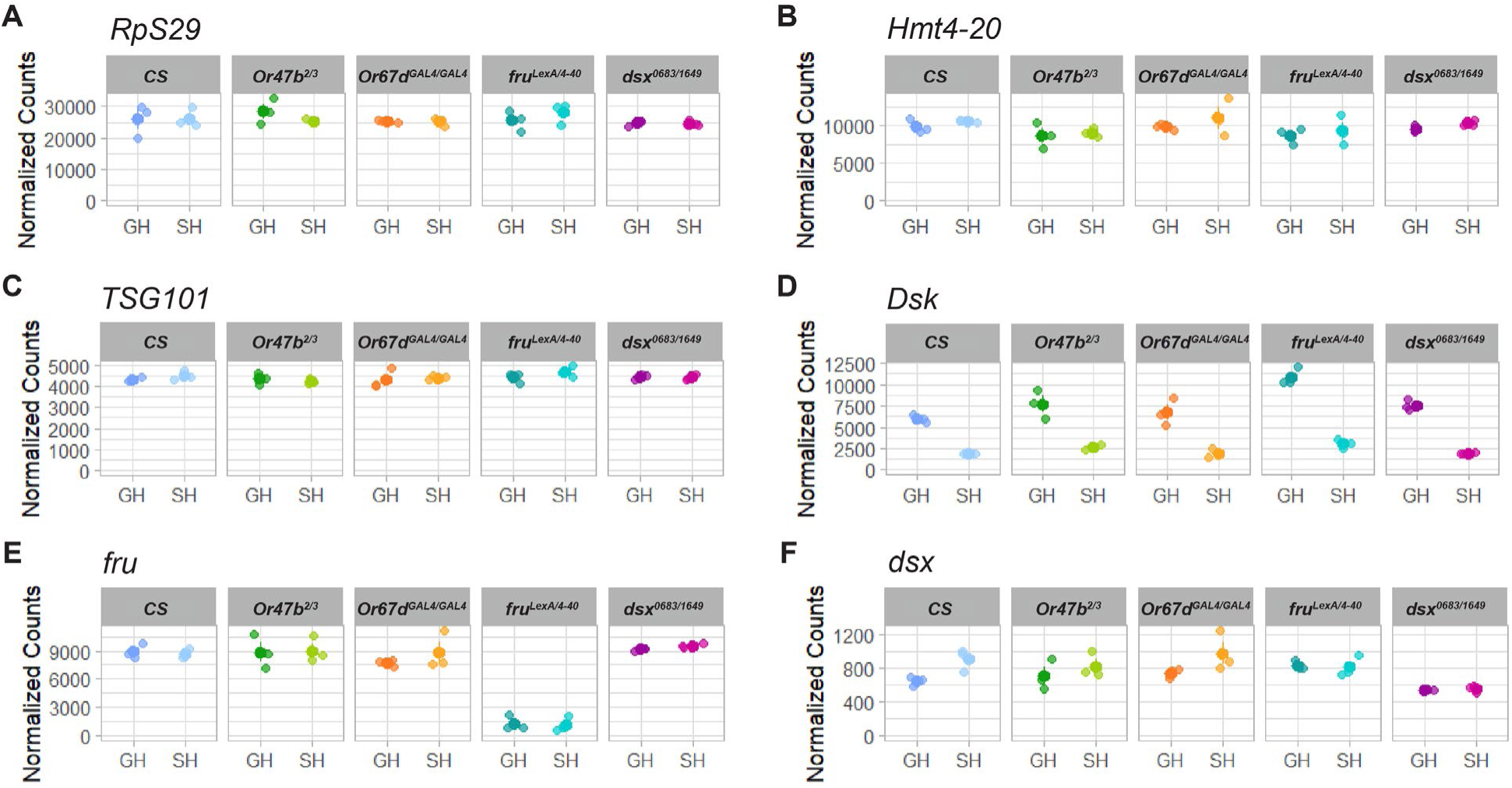
Normalized counts of genes of interest. **(A-F)** Transcript levels for several representative housekeeping genes **(A-C)**, *Dsk* **(D)**, *fru* **(E),** and *dsx* **(F)** among all wild-type and mutant brains.

**Supplementary Figure 2:**
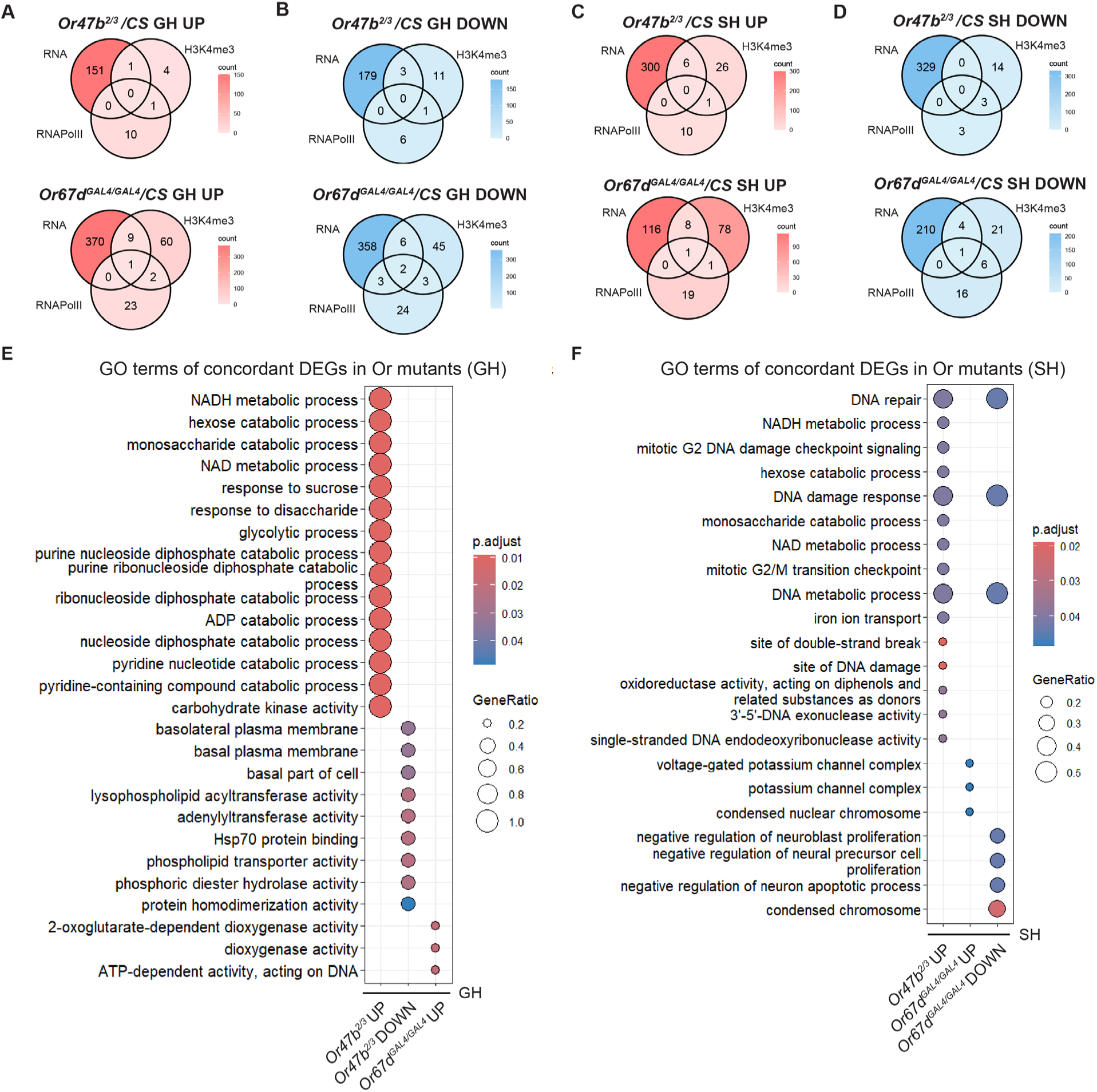
Comparisons of DEGs and DEPs in Or mutants. **(A-D)** Venn diagram comparing DEGs and genes with H3K4me3 or RNAPolII DEPs in the same direction from *Or47b^2/3^*or *Or67d^GAL4/GAL4^* mutants compared to GH **(A, B)** or SH *CS* **(C, D)**. **(E-F)** GO terms for genes showing concordant changes between transcription and at least one active chromatin mark in Or mutant/*CS* comparisons in GH **(E)** or SH **(F)** male brains (q-value < 0.05).

**Supplementary Figure 3:**
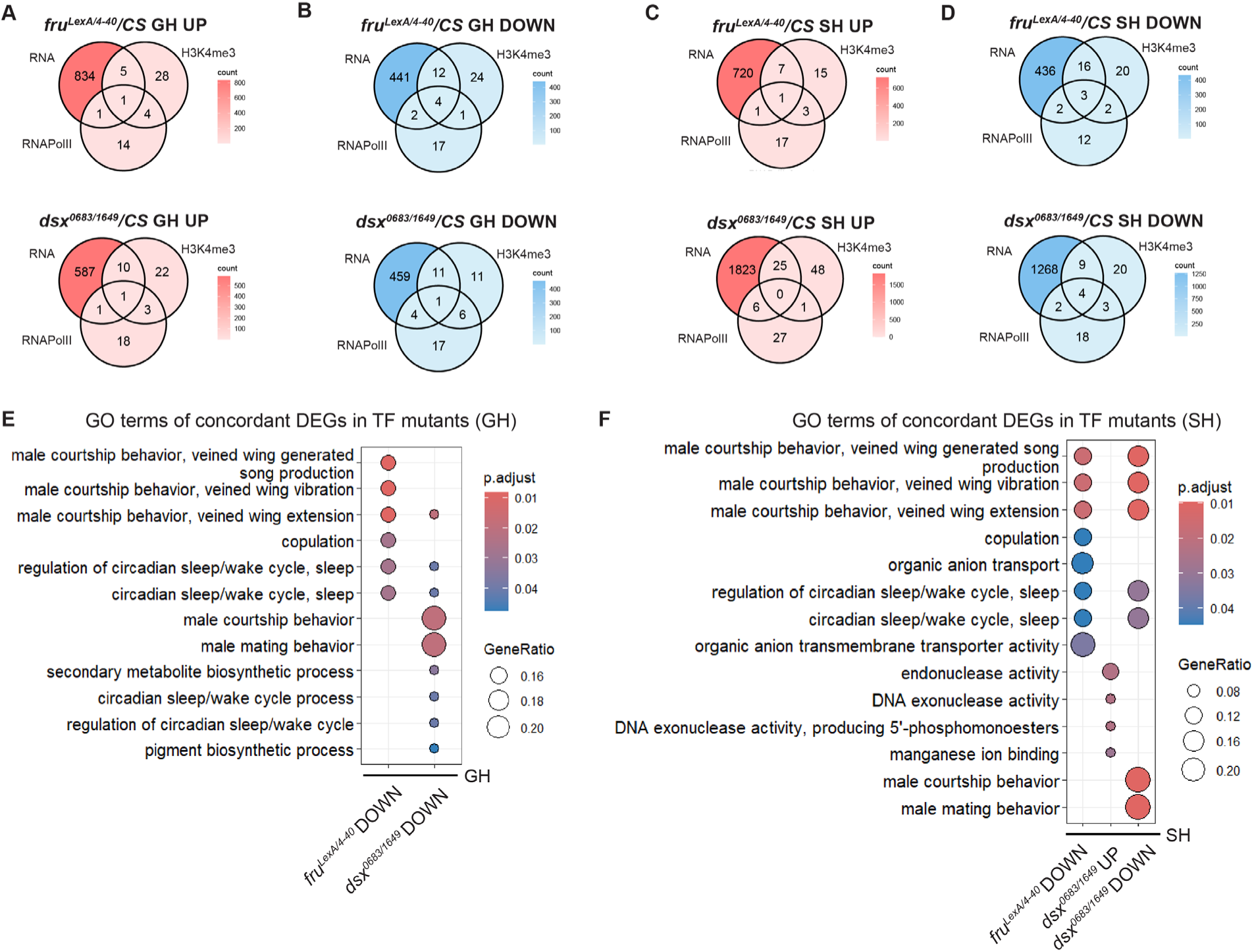
Comparisons between DEGs and DEPs in TF mutants. **(A-D)** Venn diagram comparing DEGs and genes with H3K4me3 or RNAPolII DEPs in the same direction from *fru^LexA/4-40^* or *dsx^0683/1649^* mutants compared to GH **(A, B)** or SH *CS* **(C, D)**. **(E-F)** GO terms for genes showing concordant changes between transcription and at least one active chromatin mark in TF mutant/*CS* comparisons in GH **(E)** or SH **(F)** male brains (q-value < 0.05).

**Supplementary Figure 4:**
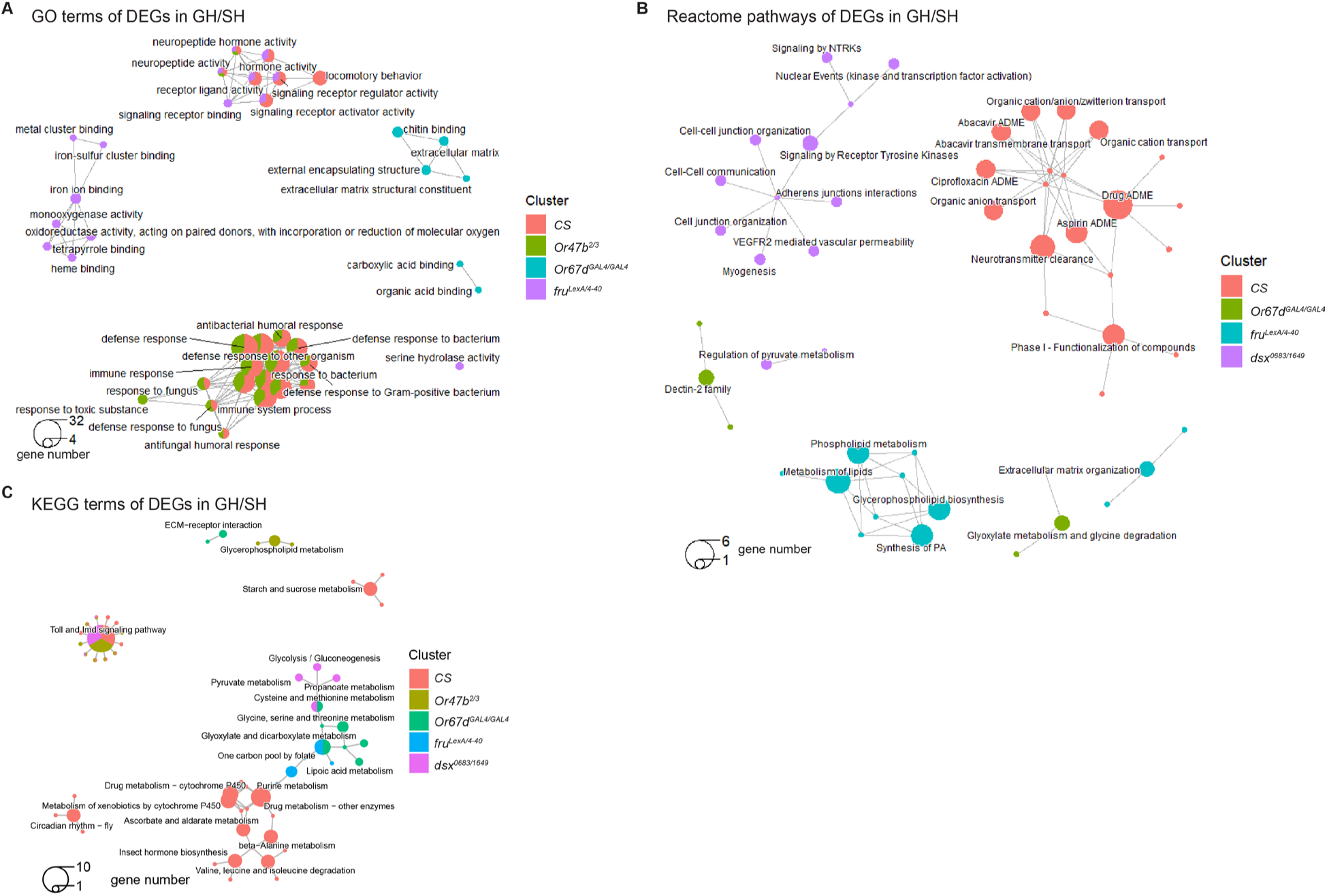
GO, Reactome, and KEGG terms of social experience dependent DEGs in *CS*, Or mutants, and TF mutants. **(A)** Connection plots of the top 15 most significantly enriched GO terms for social experience-dependent DEGs in *CS*, Or mutant, and TF mutant (q-value < 0.05) male brains. **(B)** Connection plots of the top 10 most significantly enriched Reactome pathways for DEGs from GH/SH comparisons of *CS*, Or mutants, and TF mutants (q-value < 0.05). **(C)** Connection plots of the top 10 most significantly enriched KEGG terms for social experience-dependent DEGs in *CS*, Or mutants, and TF mutants (q-value < 0.05).

**Supplementary Figure 5:**
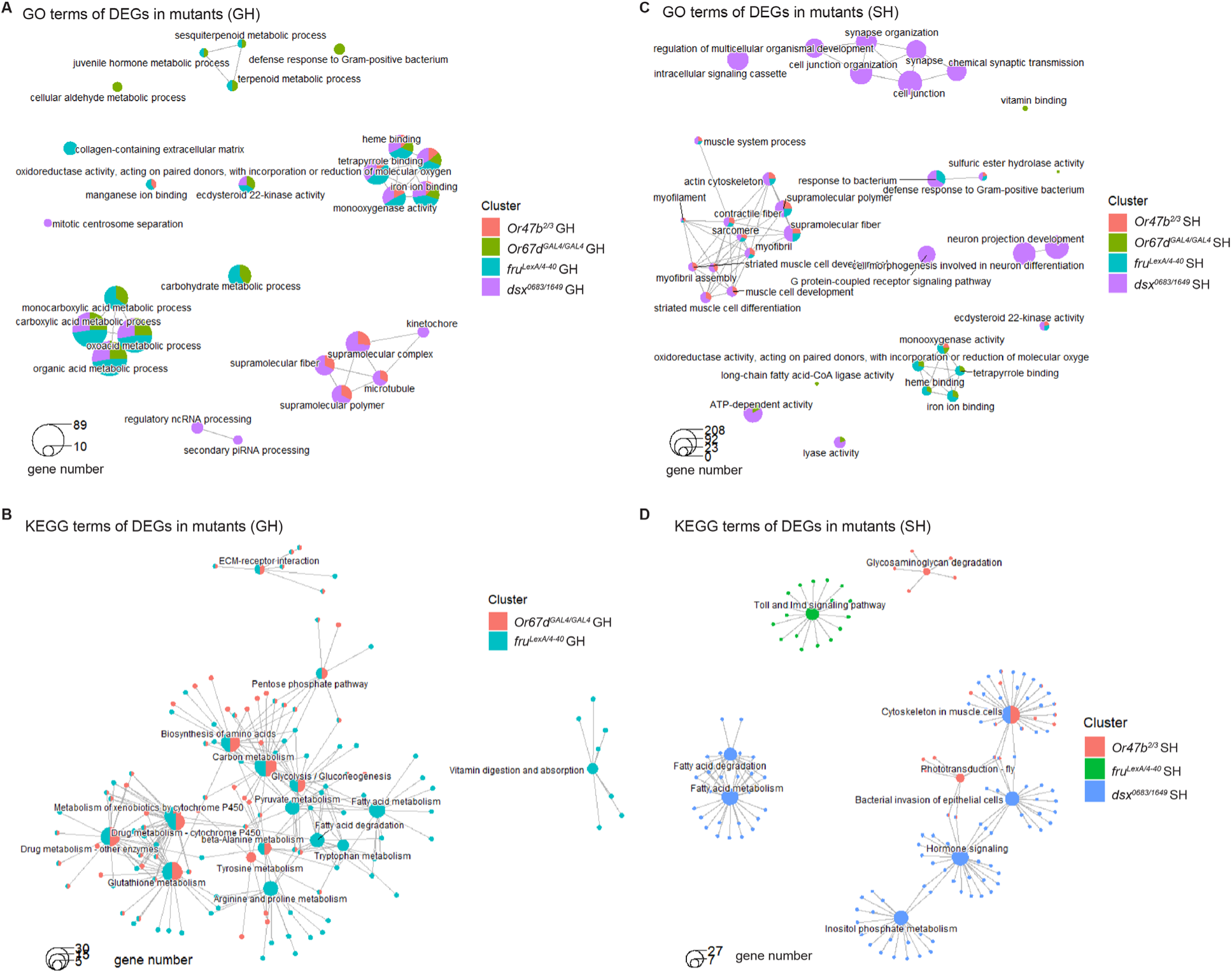
GO and KEGG terms from DEGs in mutant/*CS* comparisons. **(A, C)** Connection plots of the top 10 most significantly enriched GO terms for DEGs from mutant/*CS* comparisons under GH **(A)** or SH **(C)** conditions (q-value < 0.05). **(B, D)** Connection plots of the top 10 most significantly enriched KEGG terms for DEGs from mutant/*CS* comparisons under GH **(B)** or SH **(D)** conditions (q-value < 0.05).

**Supplementary Figure 6:**
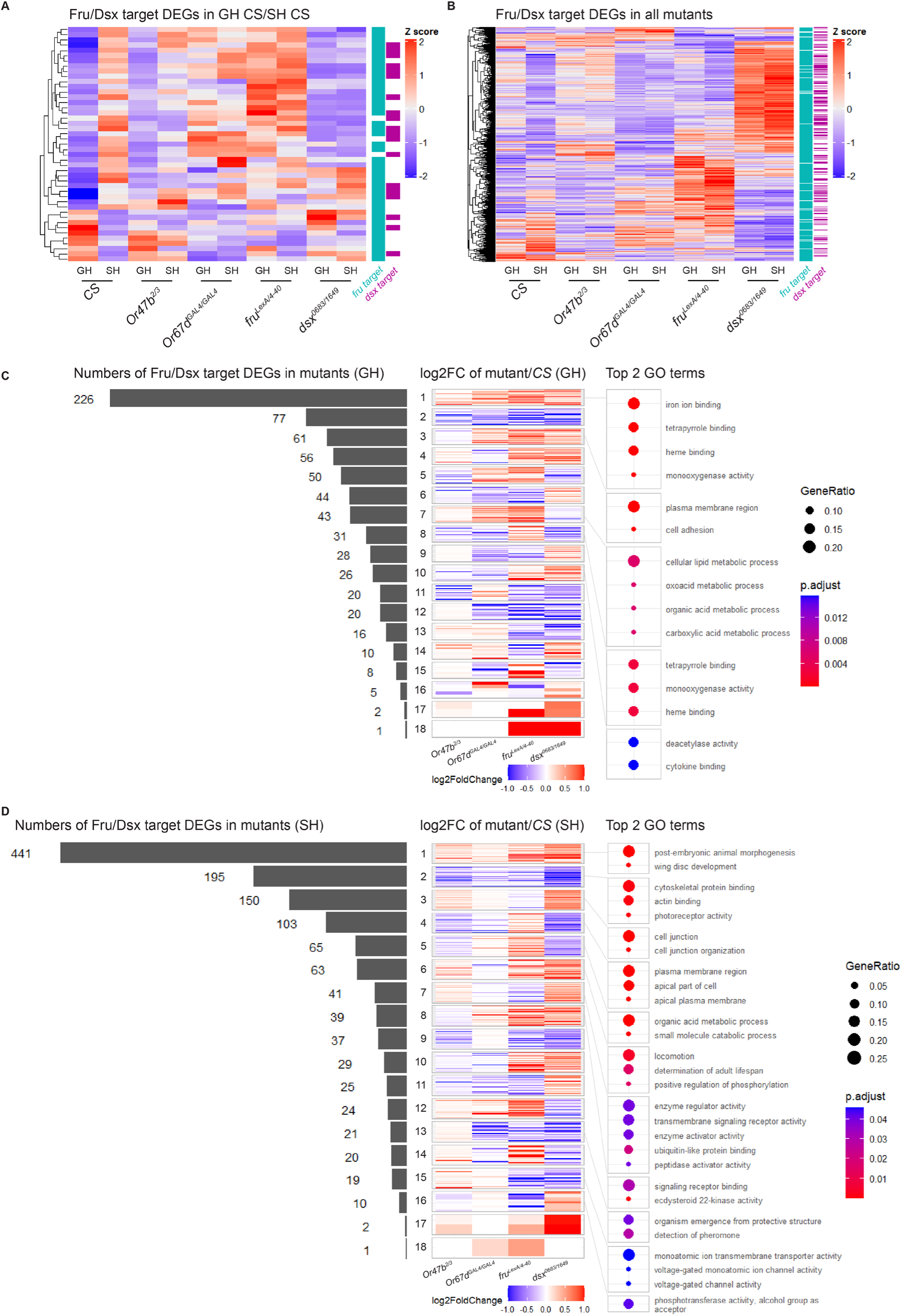
Differential effect of pheromone circuits, Fru^M^, and Dsx^M^ on social modulation of Fru^M^/Dsx^M^ target gene expression. **(A)** Heatmaps displaying Z-scores of the median of normalized transcript counts for Fru^M^ or Dsx^M^ target DEGs identified from the GH *CS*/SH *CS* comparison. **(B)** Heatmaps displaying Z-scores of the median of normalized transcript counts for Fru^M^ or Dsx^M^ target DEGs identified from the mutant/*CS* comparison under either GH or SH conditions. **(C, D)** Left: Numbers of DEGs in mutant/*CS* comparisons across various combinations of log2FC in different directions under GH **(C)** or SH **(D)** conditions. Middle: Heatmaps displaying log2FC of DEGs in mutant/*CS* comparisons across combinations of different directions. Right: Top 2 GO terms for DEGs in mutant/*CS* comparisons across combinations of log2FC in different directions in the middle.

**Supplementary Figure 7:**
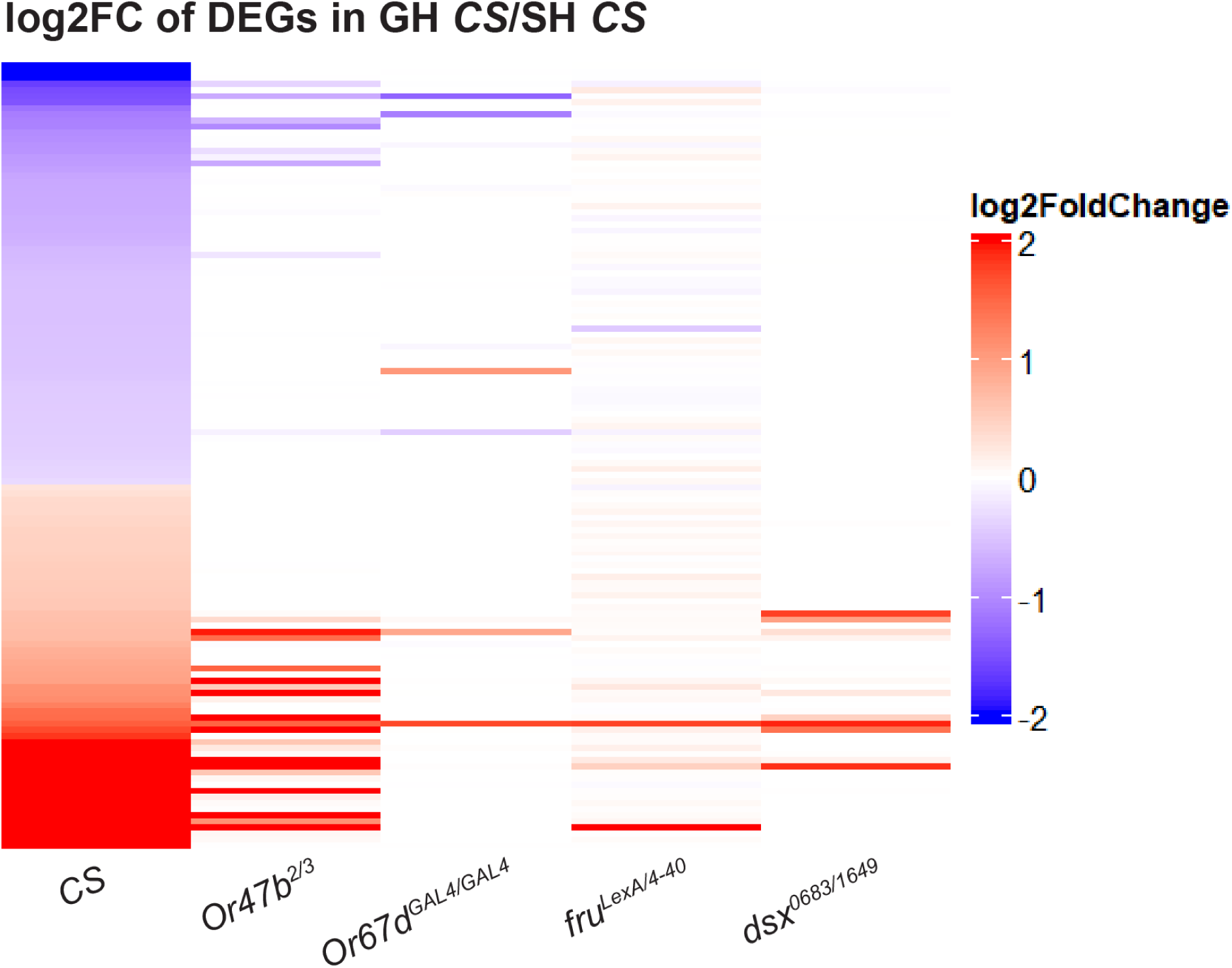
Heatmaps displaying log2FC of DEGs from GH *CS*/SH *CS* comparison in all genotypes.

**Supplementary Figure 8:**
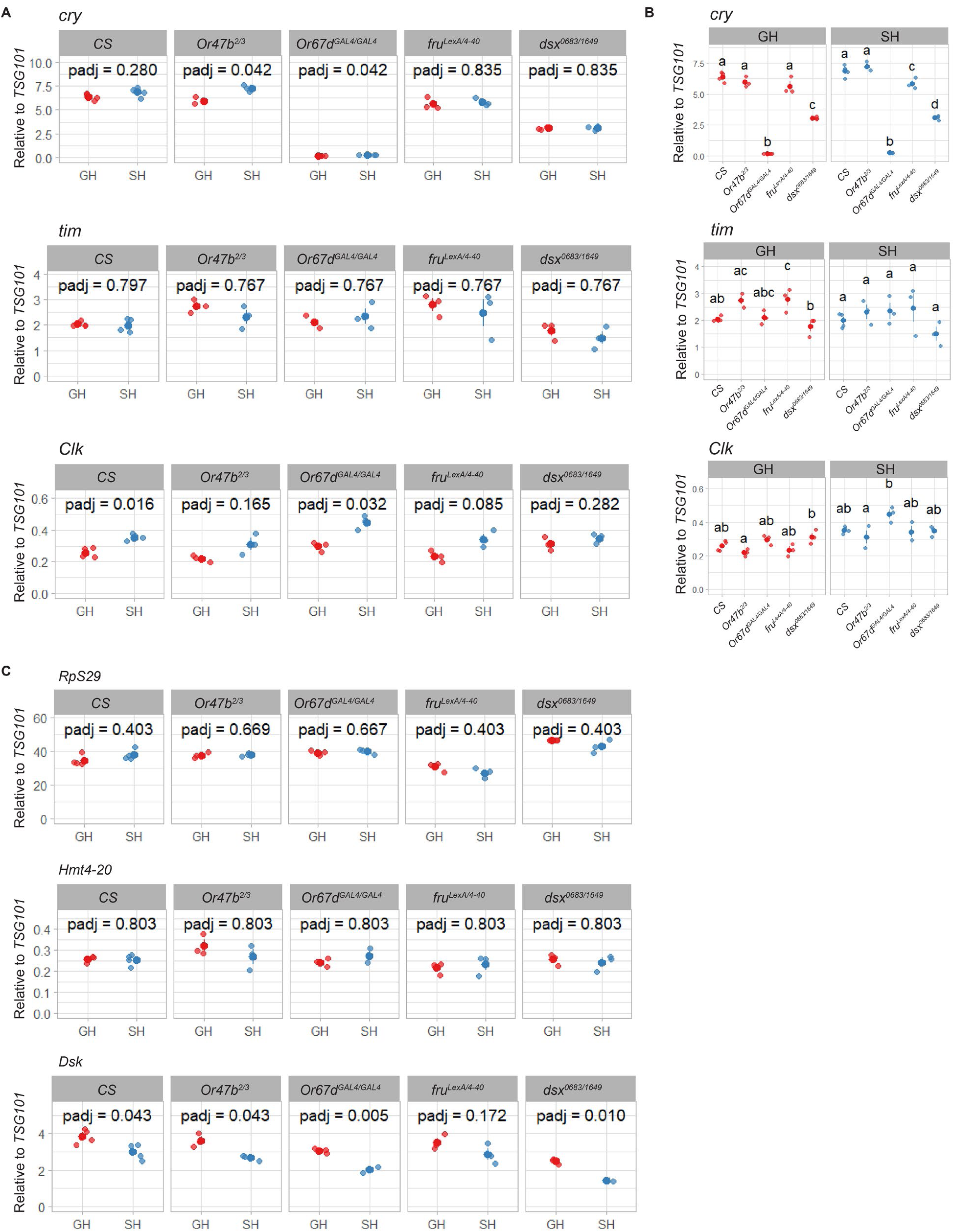
Expression of circadian genes in male heads in response to social experience. **(A)** RT-qPCR analysis of *cry*, *tim*, and *Clk* transcript levels in male fly heads between GH and SH across genotypes. Statistical comparisons were performed using two-sided t-tests. P-values were adjusted for multiple comparisons across genotypes using the Benjamini–Hochberg method. **(B)** Comparison of *cry*, *tim*, and *Clk* transcript levels in male fly heads among all genotypes under GH or SH conditions. ANOVA analysis followed by Tukey’s HSD test was applied to the statistics. **(C)** Comparison of *RpS29*, *Hmt4-20*, and *Dsk* transcript levels in male fly heads between GH and SH across genotypes. Statistical comparisons were performed using two-sided t-tests. P-values were adjusted for multiple comparisons across genotypes using the Benjamini–Hochberg method.

**Supplementary Figure 9:**
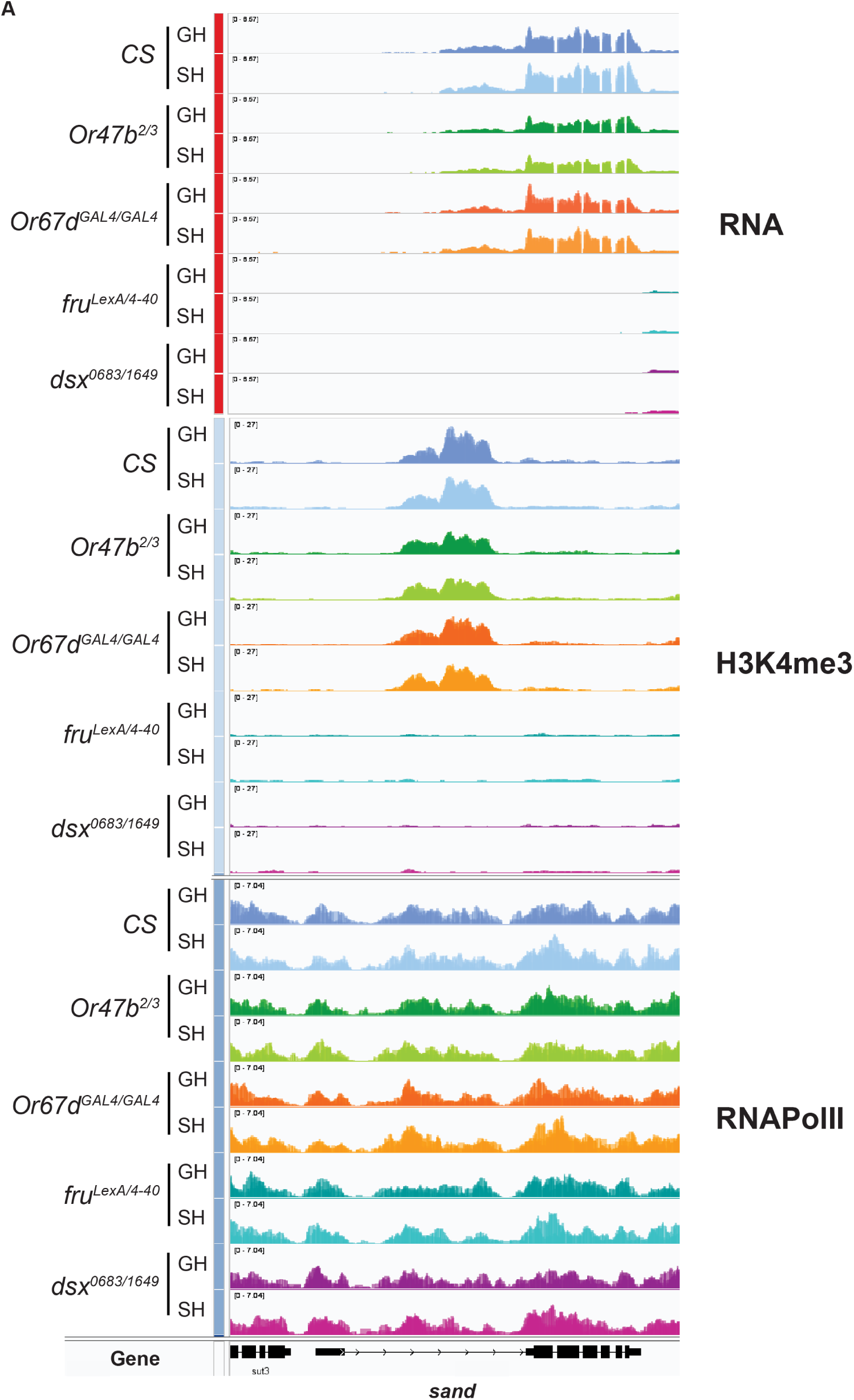
Changes in transcript and chromatin marks at the *sand* locus in different mutants and social conditions. IGV tracks showing normalized transcript levels, H3K4me3, and RNAPolII distributions at the *sand* locus across all genotypes (three biological replicates overlaid).

**Supplementary Figure 10:**
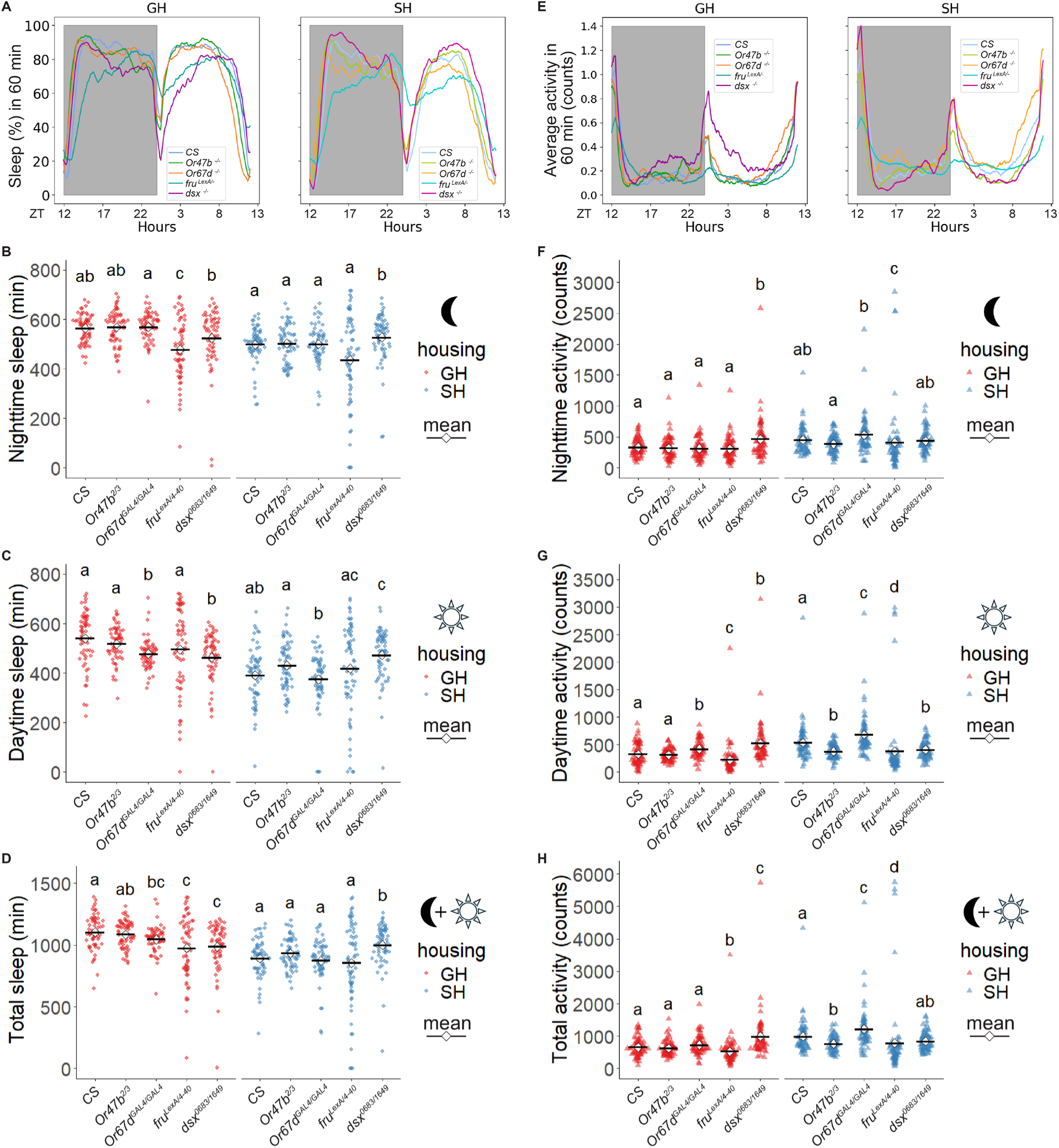
Differential effect of Or mutants and TF mutants on social modulation of behaviors. **(A)** Sleep profiles of male flies from each genotype raised under GH or SH conditions, shown as the average proportion of sleep time per 60-minute interval during a 24-hour 12:12 dark-light cycle. **(B-D)** Quantification of nighttime sleep **(B)**, daytime sleep **(C),** and daily total sleep **(D)** in **(A)**. ANOVA analysis followed by Dunn’s test was applied to the statistics. **(E)** Locomotion activity profiles of males from each genotype raised under GH or SH conditions, shown as the average counts of beam crossings per 60-minute interval during a 24-hour 12:12 dark-light cycle. **(F-H)** Quantification of nighttime activity **(F)**, daytime activity **(G),** and daily total activity **(H)** in **(E)**. ANOVA analysis followed by Dunn’s test was applied to the statistics. *P_adj<0.05, **P_adj<0.01, ***P_adj<0.001, ****P_adj<0.0001.

